# Deep Learning Algorithms Reveal Increased Social Activity of Rats at the Start of Evening Twilight

**DOI:** 10.1101/2024.07.14.603464

**Authors:** Piotr Popik, Ewelina Cyrano, Joanna Golebiowska, Natalia Malikowska-Racia, Agnieszka Potasiewicz, Agnieszka Nikiforuk

## Abstract

1.

The rapid decrease of light intensity is a potent and natural stimulus of rats’ activity. The nature of this activity, in that, the character of the social behavior and the composition of concomitant ultrasonic vocalizations (USVs) is unknown.

This study, using deep learning algorithms, sought to assess the social life of rats’ pairs kept in the semi-natural conditions at the two twilight periods. Over six days, animals were video and audio recorded during the morning and the evening sessions lasting for 20-minutes each. The videos were used to train and use the DeepLabCut neural network examining animals’ movement in space and time. Numerical data generated by DeepLabCut were subjected to the Simple Behavioral Analysis (SimBA) toolkit, to build models of 11 distinct social and non-social behaviors. DeepSqueak toolkit was used to examine USVs.

Deep learning algorithms revealed lights-off induced increases of fighting, mounting, crawling, and rearing behaviors, as well as of 22-kHz alarm calls and 50-kHz flat and short, but not frequency modulated calls. In contrast, lights-on stimulus increased the general activity, as well as adjacent lying (huddling), anogenital sniffing and rearing behaviors. The animals adapted to the housing conditions showing decreased ultrasonic calls, as well as of grooming and rearing behaviors, but not fighting.

Present study shows lights-off induced increase of aggressive behavior but fail to demonstrate an increase in the positive affect defined by the hedonic USVs. We further confirm and extend the utility of deep learning algorithms in analyzing rat social behavior and ultrasonic vocalizations.

**Highlights:**

- Rats display a natural increase in activity induced by lights-off stimulus
- Deep learning algorithms allow for rapid characterization of social behavior and ultrasonic calls
- The darkness increased aggressive behavior, 22-kHz alarm calls and 50-kHz flat and short, but not frequency modulated calls

## 2. Introduction

Every researcher who has ever entered the animal room in the evening, has noted rapid increase of rats’ activity. Wild rats are nocturnal or crepuscular, and most of their activities occur under low-light conditions [1, 2]. The laboratory rats retain this propensity [2, 3]. This observation has several consequences. For instance, it implies the necessity of mimicking the natural light-dark cycle in the animal facilities, and the importance of studying rats’ behavior at their active (dark) phase. As rats exhibit the positive and negative affect, expressed as the 50-kHz “happy” and 22-kHz “alarm” ultrasonic calls, respectively (see [4] for the recent review), it is also likely that the “positive” affect is more intense at the active (dark), than at the resting (light) phase. In this respect, Burgdorf et al., [5] reported that the lights-off is a strong signal for the induction of locomotor activity and pro-social behavior resulting in hedonic 50-kHz ultrasonic vocalizations (USVs), [5].

The rapid decrease of light intensity, signaling the active rats’ phase appears as a perfect natural phenomenon for studying the motivation, because as the animals wake up, they begin their social and other activities. Here we attempted to measure rats’ general activity, social behavior, and USVs using objective and top modern digital techniques to find whether these can “catch”, and further elucidate, the nature of darkness-induced activity in semi-natural conditions.

To this aim, we have built 6 custom sound-attenuated boxes equipped with the lickometers monitoring head entries into the water supply, infrared (IR) sensitive cameras registering social activities, and the microphones that recorded animals’ ultrasonic calls. The boxes were also equipped with electric fans supplying fresh air, white lights controlled by an ON-OFF clock, as well as with the IR LEDs providing conditions necessary for video recording at the total darkness.

This setup was designed to precisely, automatically and objectively monitor rats’ activity at the two twilight periods, i.e., when the lights rapidly were turned off in the evening, and on in the morning sessions, respectively. Prompted by Burgdorf et al., [5] research, we expected high intensity of prosocial activity and of the positive affect at the beginning of the dark phase. Studying these phenomena at the end of the dark phase was regarded as control conditions.

Rat’s social life [6] is extremely complex [1, 7, 8]. Thus far, the rodent social behaviors and ultrasonic vocalizations have been classified by the trained observers who scored them either at the time the tests were conducted, or off-line, by watching the video tapes or computerized videos, and analyzing and classifying USVs manually on a computer screen (see our recent paper by Popik et al., [9] for more details). Such approach, however, is extremely time-consuming and prone to the subjective bias.

Recent advances in computer vision and deep learning-based toolkits enabled almost semi-automatic video analysis of the rodent social behavior [10]. Specifically, behavioral analyses offered by the open source DeepLabCut https://github.com/DeepLabCut marker-less pose-estimation toolkit [11–13] have vastly facilitated the analytical workflow. This is because DeepLabCut offers the generation of neural networks (models) representing interacting animals’ body parts in time and space. This freely available software allows for semi-automatic tracking of animals’ body parts movements. A necessary second step requires post-processing software, that using numerical data representing animals’ body parts in time and space (i.e., the output provided by the DeepLabCut), would classify social behavior. For these, the Simple Behavioral Analysis (SimBA, https://github.com/sgoldenlab/simba) open source python’s toolkit, constitutes another analytical break-through [14, 15]. The third digital tool used in this work was the DeepSqueak https://github.com/DrCoffey/DeepSqueak, the Matlab’s machine learning software that vastly facilitated the analysis of ultrasonic calls [16].

This work has focused on examining whether darkness-induced activity could be detected with the use of semi-automatic and objective digital workflow and, on finding the precise nature of the activity, expressed by the analysis of social behavior and USVs categories. The availability of categorized ultrasonic and behavioral data allowed also for investigation of the relationships between them.

## 3. Methods

### 3. A. Animals and Ethics

Twelve male 7-week-old Sprague Dawley rats (Charles River, Germany) weighing ∼ 300 g upon arrival were group housed (4 per cage) in the standard laboratory cages for 2 weeks of acclimation. Both at that time, as well as during the experiment, the animals were maintained under standard colony A/C controlled conditions: room temperature 21 ± 2 °C, humidity (40–50%), 12-hr light/dark cycle (lights on: 06:00) with *<u>ad libitum</u>* access to water and lab chow.

The animals were maintained, and experiments were conducted in accordance with the European Guidelines for animal welfare (2010/63/EU) and EQIPD guidelines. All experimental procedures were approved by the II Local Ethics Committee for Animal Experiments at the Maj Institute of Pharmacology, Polish Academy of Science, Kraków, Poland (ethical allowance LKE: 108/2023).

### 3. B. Apparatus

The experiment was conducted in six identical custom boxes (length x width x height: 50 x 50 x 50 cm) made of the black Plexiglas. Every box’ top contained: a) 20 holes (diameter of 2.5 cm) allowing for fresh air circulation, b) a hole of 7.5 cm diameter with 12 V electric fan (Sunon, China) facilitating airflow, c) infrared (IR) sensitive Axis M1137 Mk II network camera (Axis Communications, Sweden) connected via Ethernet ports to the Synology NAS station, d) Avisoft UltraSoundGate CM16/CMPA ultrasonic microphone connected via UltrasoundGate 416H analog-digital converter into one of two laptop computers running RECORDER USGH software (version 4.2; Avisoft Bioacoustics; Glienicke/Nordbahn, Germany), e) white light LED strip providing 150 Lux (measured at the box’s bottom) of ambient white light, controlled by a precise circadian clock, and, f) two IR LEDs providing the illumination necessary for video recording at the total darkness. The box’s front had the 4 x 5.5 cm (width x height) opening located 7 cm above the floor, handling the water bottle + sipper tube (Med Associates model ENV-250BT) water supply, monitored with the IR Head Entry Detector (Med Associates model ENV-254-CB) connected via Med Associates SG-716B SmartCtrl™ Interface Modules and DIG-705 box to the third laptop computer. The inner surfaces of the boxes were covered with the P80 K20N (Norton, Poland) sandpaper up to the height of 24 cm; the rest of the inner walls were covered with the 2-cm thick black sponge (BitMat, Poland) restricting echo and allowing for good quality USV recordings. The boxes’ bottoms were covered with the Multifit [J1230123], Germany) Forest-Land wood shavings, allowing for a dark, contrasting background. The lab chow was freely available at the boxes’ floor. All 6 boxes were placed in the same animal colony room, dedicated to this experiment.

### 3. C. Procedure and Experimental design

Two unfamiliar rats of matched body weight (± 5 g) were placed in the custom box and remained there undisturbed for 6 days, i.e., until the end of experiment. The water supply head entries were monitored continuously, while the video and audio recordings were done for 20 minutes daily, in the morning sessions (05:50-06:10) and for 20 minutes in the evening sessions (17:50-18:10). The videos (mp4 H.264 encoded, 640 x 480 pixels, variable bitrate, image quality 3), were recorded at 12 frames per second (fps), while audio data were recorded at 250,000 Hz.

As this work should be regarded as exploratory, we did not perform sample size analysis. As in this work we did not investigate differences among groups, the experimental protocol had no randomization and blinding.

### 3. D. Digital workflow

We provide a detailed workflow allowing easy reproduction of the steps leading to the semi-automated analysis of behavioral and USV data. While the DeepLabCut https://github.com/DeepLabCut , SimBA https://github.com/sgoldenlab/simba and DeepSqueak https://github.com/DrCoffey/DeepSqueak websites offer comprehensive details of the installation and use of the respective toolkits, we describe our workflow as an easy step-by-step guide for the less computer-oriented researchers.

### 3. E. DeepLabCut frames’ labeling, training, and network evaluation

We have chosen 37 random videos (**Figure** 1) and labeled 582 random PNGs with 12 body parts of rat A and 12 body parts of rat B (nose, left eye, right eye, head, back, pelvis, anogenital area, left shoulder, right shoulder, middle, tail middle, and tail end).

**Figure 1.**
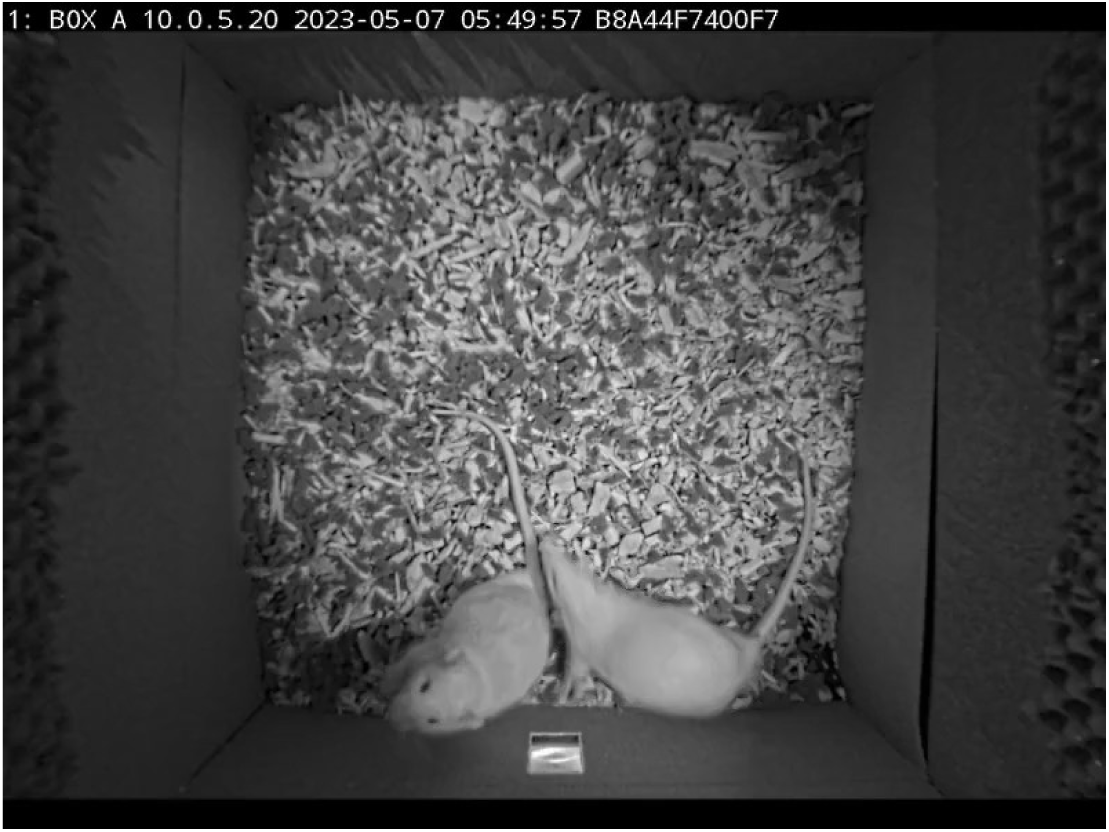
A raw video analyzed by DLC shows the camera view with two white rats, a water supply at the bottom, a sandpaper and echo-attenuating sponge on the inner walls.

Care was taken to aim for as much of variety in behavior, posture, individuals used, and backgrounds, as it was possible within the dataset, as suggested by Hardin and Schlupp, [17].

Labeling was done over several days using DeepLabCut “napari” plugin on a desktop PC with Windows 11 equipped with the Nvidia RTX 4090 graphic card and the GUI version 2.3.5 of DeepLabCut running on Python 3.8.16.

To achieve the best possible model, we trained several DeepLabCut “shuffles”, gradually increasing the number of annotated frames, eliminating badly recognized frames, and varying the number of iterations. We finally trained DeepLabCut model with 200,000 iterations. Other DeepLabCut’s variables were set at default values: default_net_type: dlcrnet_ms5, default_augmenter: multi-animal-imgaug and default_track_method: ellipse. This was done iteratively over several weeks on the Nvidia DGX A100 station running Ubuntu 22 Linux, Python 3.9.15 (main, Nov 4 2022, 16:13:54), IPython 8.6.0, and DeepLabCut version 2.3rc2.

As with other variants, also the shuffle #37 and its snapshot index “-4” (170,000 iterations) was followed by network evaluation (**Figure** 2). Shuffle #37 was chosen as a final model because it displayed the lowest Train error and Test error of 3.45 and 5.96 pixels, respectively with p-cutoff of 0.6. As a **check point**, the videos that were analyzed with shuffle #37 were played on a computer screen at reduced speed with the free mpv program https://mpv.io/. We noted that DeepLabCut marked the body parts the way the observer would mark them, and that the incidents of interchanging body part(s) between rats, or of “loosing” the animal from the view, were sporadic.

**Figure 2.**
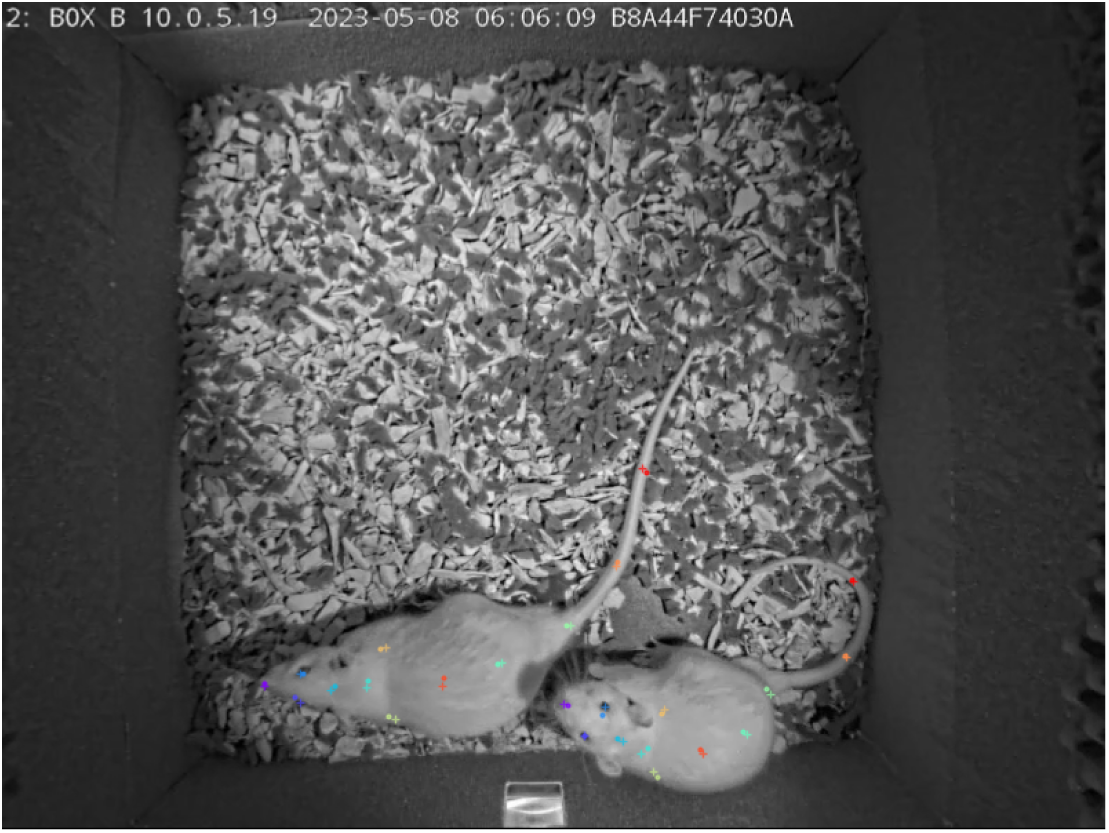
A frame showing human annotations (+) and how accurately DLC recognized them (.)

As a result, DeepLabCut analysis provided the videos with the “skeletons” and the numerical data (CSV files) representing every body part position in space and time; all these files were used by SimBA toolkit in the next steps.

### 3. F. Post-DeepLabCut processing with SimBA

Using shuffle #37 we analyzed 16 SimBA training videos, randomly chosen from the 39 videos of the morning sessions and 33 videos of the evening sessions. We ran SimBA ver. 1.87 GUI, loaded the initialization file, and imported CSVs with INTERPOLATION METHOD: Body Parts Nearest, and SMOOTHING: NONE. We skipped OUTLIER CORRECTION.

### 3. G. Extracting SimBA’s “features” with two custom scripts

In the next step, we processed the input CSVs using python 3.6 scripts to save the custom set of “features”, which are a complex assembly of settings (e.g., distances, movements, etc.), used to recognize various relationships between body parts. Since this approach differs from the default SimBA’s workflow, it is described in more detail.

Our initial attempts with the use of a default set of features offered by a standard SimBA’s workflow failed to provide decent classifiers. SimBA offers to use additional set of features, including, among others, the angles, convex hulls etc. However, even the enlarged set of features did not provide well-performing classifiers, and their analyses have often hung the program due to computer’s memory limits. For these reasons, and prompted by Lapp et al., work [18], we decided to create a custom set of restricted features of interest.

The CSVs located in SimBA’s “outlier corrected movement location” folder were first processed by the two custom scripts.

### 3. H. Dr. Simon Nilsson’s polygons script

The script^1^, kindly provided by Dr. Simon Nilsson and slightly modified, produced the “polygons” features (“polygon_pct_overlap” and “difference_area”). These two features served mainly to detect how close the animals were to each other, see **Figure** 3. However, the script also filled the missing body parts using data_df.interpolate(method=’nearest’).bfill().ffill() python panda’s function, which “guessed”, where possible, the position of all body parts and filled them appropriately. Such interpolation of body parts positions was necessary, because the random forest machine learning approach requires all cells not to be empty. The missing body parts were filled with 0 using another script^2^.

**Figure 3.**
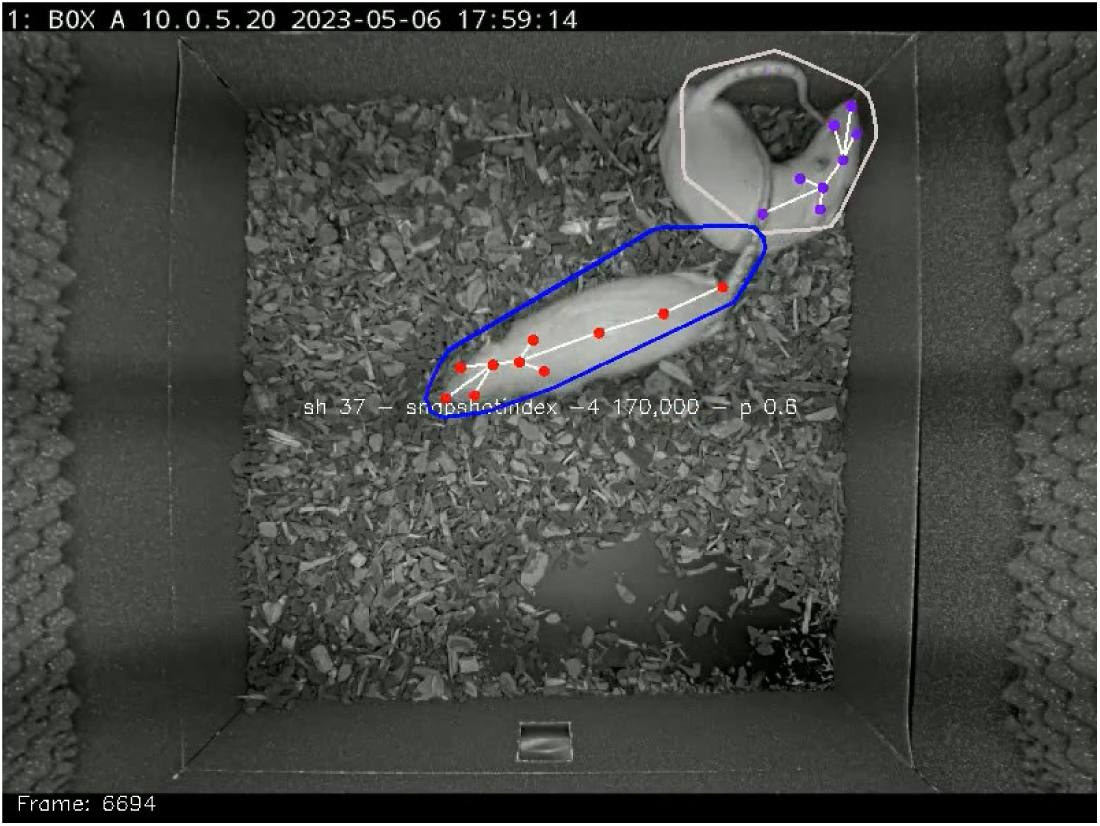
Dr. Simon Nilsson’s polygon features in action

### 3. I. The custom features script based on Dr. Hanna Lapp’s “Amber” project

Script^3^ was based on Lapp et al., work [18], fully described at the github site https://github.com/lapphe/AMBER-pipeline/blob/main/AMBER_pose_estimation.py

We modified it to suit present experimental conditions: that is, to analyze relationships of two similarly looking rats, rather than of a dam and her offspring. The script calculated several features, including the whole body, head and anogenital area “centroids” that is x, y coordinates indicating the center of a given body part.

The genuine advantage of analyzing the centroids (rather than solely individual body parts’ points) is that even if a given body part was temporarily missing due to the occlusion by other animal, the centroids were almost always detected and could serve for calculating other features.

These derived features included the distances between different animals, for example, the heads (likely reflecting nose contacts and sniffing behavior) or the distances between the head and anogenital region (likely indicative of anogenital sniffing). Other behaviors’ detection, like mounting, crawling, nosing, and following also depended on the distances between centroids of different animals. The script calculated also distances between same animal body parts (useful for determining rearing and self-grooming behaviors).

While the centroids were saved as the pairs of x, y coordinates, functionally similar measures were the convex hulls, presenting the area of the animals’ whole body (likely useful for detecting rearing behavior), and the head (which could change its shape when the nose of the animal was not visible due to sniffing or self-grooming). For the same reason we programmed the script to determine the movements including 1-second rolling movements (likely useful for detecting the following behavior), as well as the angles between specific body parts (likely helpful to identify rearing behavior). Finally, the script saved the probabilities of detections as this helped to determine behaviors that were less likely to be detected (as e.g., during fast moving or in cases one animal occluded another when mounting or crawling). In total, the script determined 109 custom features.

Script^4^ allowed for visualization of the features saved in the previous step (**Figure** 4). As a **check point**, the videos were inspected at the slow motion.

**Figure 4.**
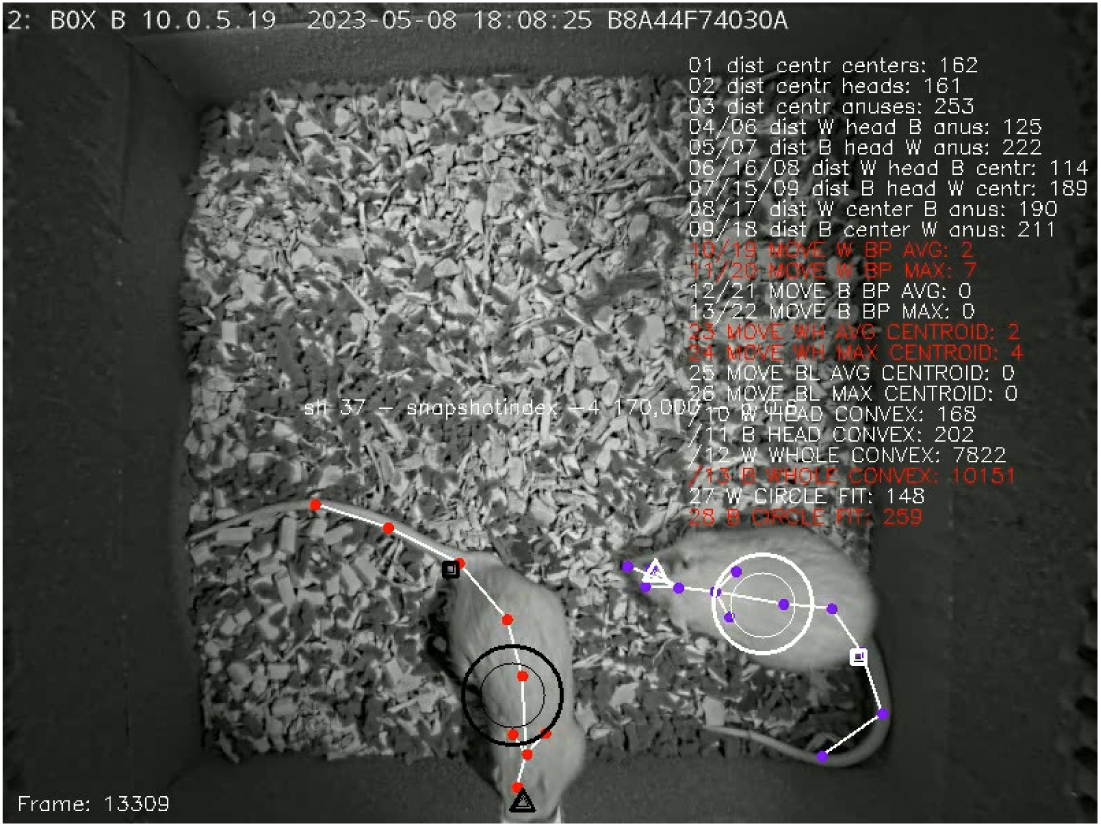
Dr. Hannah Lapp’s - like custom features in action. The circles, triangles and rectangles correspond to the whole body, head and anogenital “centroids”, respectively. The larger elements represent the actual position. The smaller elements represent 1-second rolling mean position. Red text indicates that a given feature was active, for instance, given body parts were closer than the threshold, or in movement.

### 3. J. Merging the polygons with Amber-like features into one set

Unnecessary columns were removed from the CSVs using the script^5^ so that only 109 columns with the features were left. Then, Dr. Simon Nilsson’s 2 polygon features CSVs and Dr. Hannah Lapp’s-like 109 features CSVs were merged using another script^6^ that also adjusted the number of rows in every CSV to the minimal number. This created the final “features extracted” set of CSVs, which was copied to the “features extracted” folder.

### 3. K. SimBA training

We analyzed the following 11 behavioral categories: 1 **adjacent lying** (the time of side by side contact or “huddling”; see [19, 20]), 2 **anogenital sniffing** (one rat sniffing the anogenital region of the conspecific), 3 **crawling** (one rat moving over or under the conspecific), 4 **fighting** (mostly aggressive grooming: one rat chasing another then, while pinning down a conspecific or holding it with the forepaws: licking, chewing the fur of the conspecific, or punching; see [14, 15, 21–23]), 5 **following** (active movement of two individuals; one chasing and approaching another), 6 **grooming** (known also as allogrooming: cleaning the fur or amicably scratching, licking and chewing the fur of the conspecific, motionless [22]), 7 **mounting** (climbing or standing on the back of the conspecific), 8 **nosing** (rats touching each other with their noses, while stretching their body slightly; [24]), 9 **rearing** (one or both animals standing on their hind legs), 10 **self-grooming** (cleaning the fur or scratching: rapid movements of the head towards the own body; [24]) and 11 **sniffing** (sniffing or touching the body of the conspecific). The choice of these behaviors was based on a number of ethological observations [1, 7, 8, 14, 15, 21–24] and represents highly characteristic set of behaviors of the same-sex *laboratory* rats in dyadic encounters. Representative examples of the 11 scored behavioral categories are shown on **Figure 5**.

**Figure 5.**
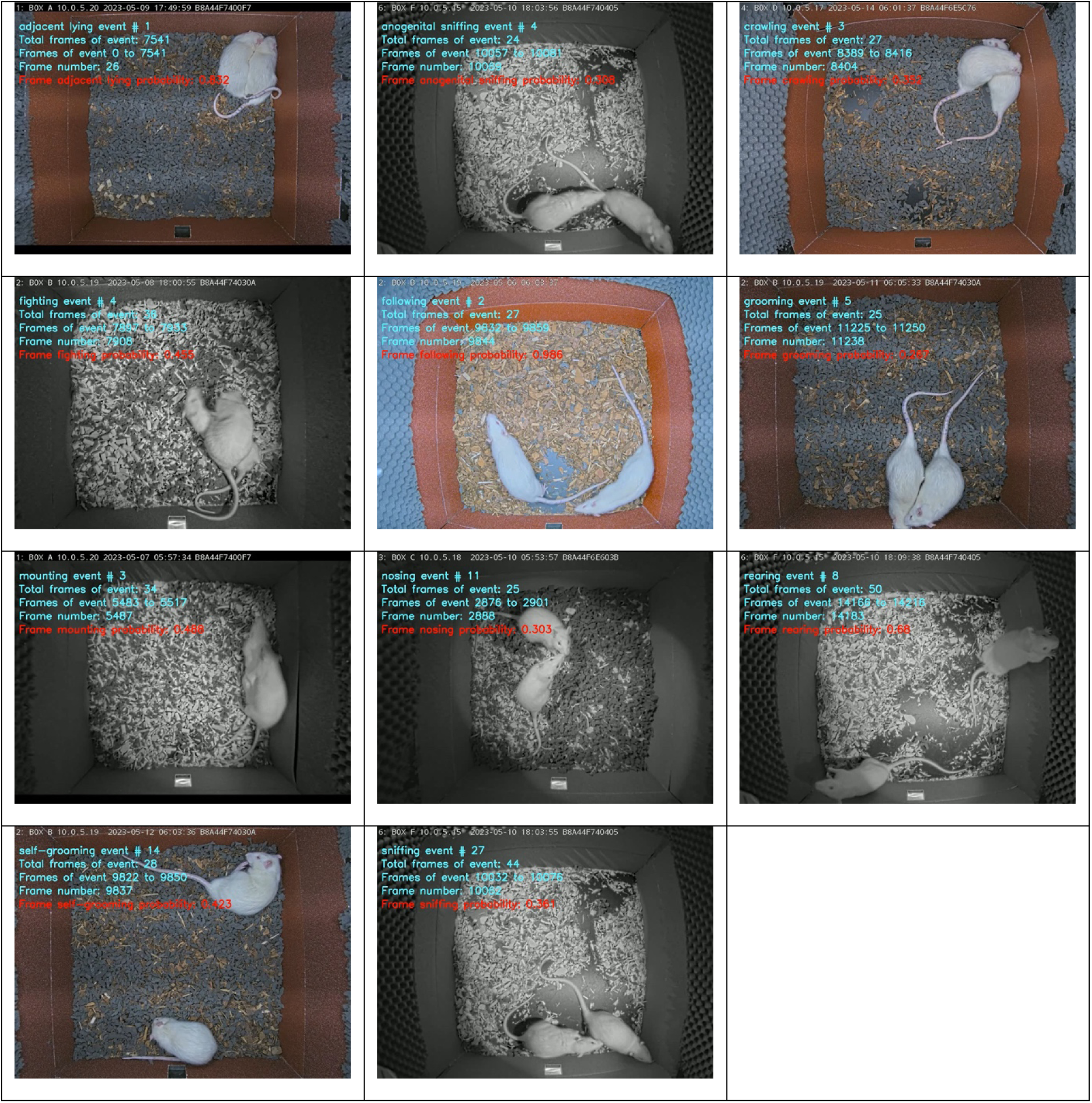
Representative frames showing 11 behavioral categories.

Having a custom set of 2+109 features, it was possible to train SimBA models (classifiers). We used 16 randomly chosen videos, some of a short duration but rich in a given social behavior. As we labeled frames with the behaviors earlier, using the default set of features, the CSVs columns with individual classifiers displaying 1 (behavior present) or 0 (behavior absent) and stored in the “targets inserted” training folder were combined with the CSVs stored in “features extracted” folder using the script^7^.

Following training, the examination of classifiers’ performance using summary precision curves (**Figure** 6) with the custom script^8^ revealed decent F1’s and allowed for setting the initial Detection Thresholds, see **Table 1**.

**Figure 6.**
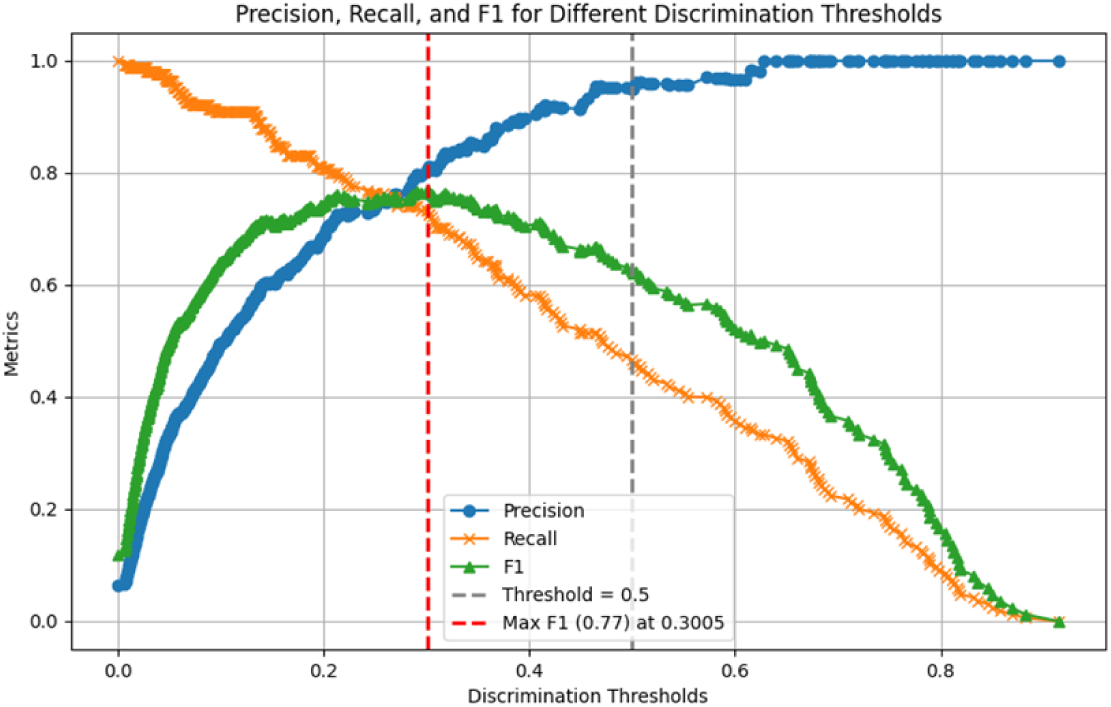
A sample precision curve of “anogenital sniffing” classifier shows its maximum F1 = 0.77 at Detection Threshold ∼ 0.3. This Detection Threshold was initially set in the SimBA’s INI file to examine this classifier’s performance.

**Table 1.**
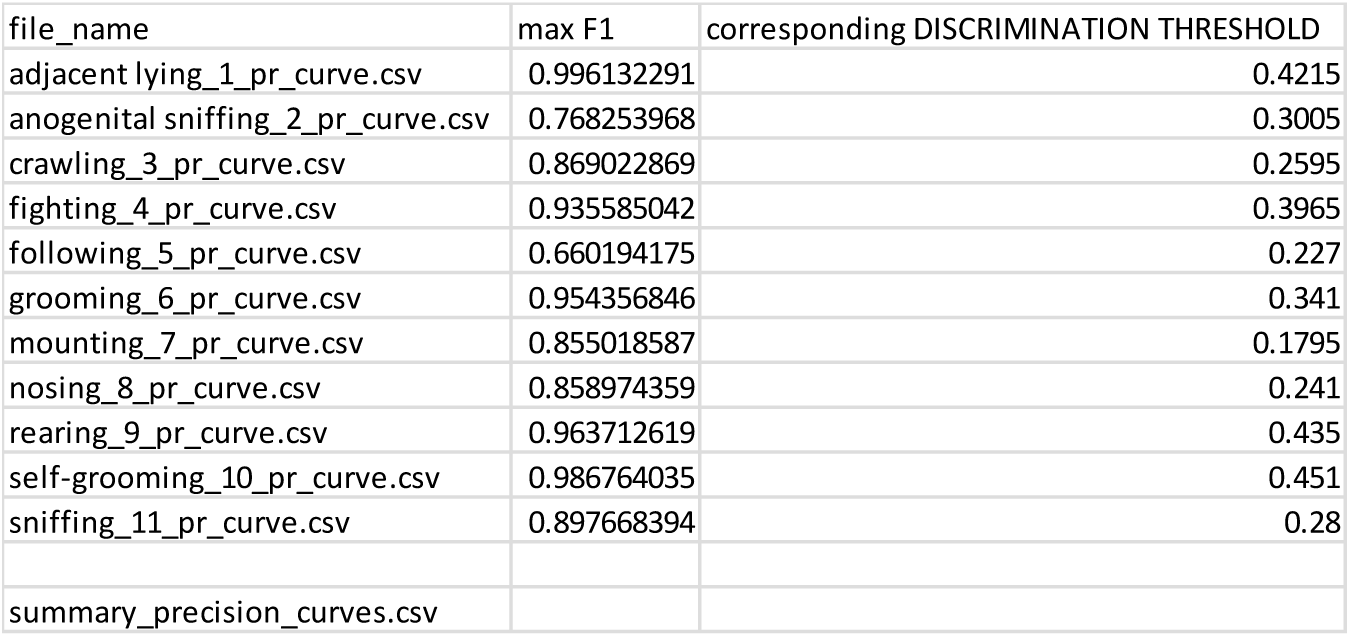
The performance (F1) of 11 behavioral classifiers (models) and their corresponding Detection Thresholds.

| file_name | max F1 | corresponding DISCRIMINATION THRESHOLD |
| --- | --- | --- |
| adjacent lying_1_pr_curve.csv | 0.996132291 | 0.4215 |
| anogenital sniffing_2_pr_curve.csv | 0.768253968 | 0.3005 |
| crawling_3_pr_curve.csv | 0.869022869 | 0.2595 |
| fighting_4_pr_curve.csv | 0.935585042 | 0.3965 |
| following_5_pr_curve.csv | 0.660194175 | 0.227 |
| grooming_6_pr_curve.csv | 0.954356846 | 0.341 |
| mounting_7_pr_curve.csv | 0.855018587 | 0.1795 |
| nosing_8_pr_curve.csv | 0.858974359 | 0.241 |
| rearing_9_pr_curve.csv | 0.963712619 | 0.435 |
| self-grooming_10_pr_curve.csv | 0.986764035 | 0.451 |
| sniffing_11_pr_curve.csv | 0.897668394 | 0.28 |
| summary_precision_curves.csv |  |  |

Every classifier was then iteratively assigned with its detection threshold (how sure the program must be, to classify a given frame as containing a given behavior). These values were tweaked by inspecting the resulting videos to provide even better detections. Each classifier has been assigned with its minimal bout length = 0 so if present, it was represented in the “frame”, and was not required to last for a longer time than 1/12 s. This allowed for the precise assessment of behaviors’ duration, but not for the number of given behavioral category episodes (see **Table** 2 below).

The classifiers were created using the random forest approach, entropy criterion, 20% of the training data, and no under/over sample parameters.

### 3. L. SimBA testing

Having well-performing classifiers, the testing videos were examined for the specific behaviors. The script^9^ was run to store annotated videos in a custom folder. As a **check point**, we examined ensemble predictions in a custom way to observe how the classifiers detected the behaviors (**Figure** 7).

**Figure 7.**
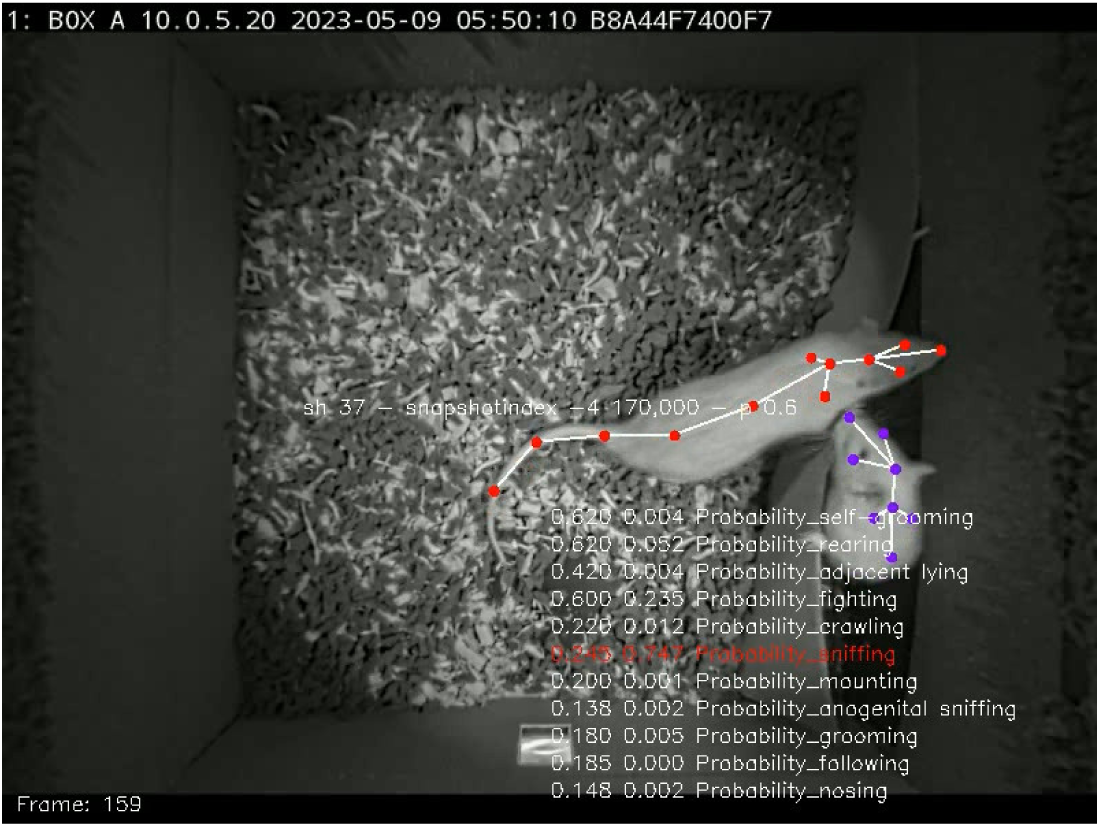
A frame from the video with behaviors annotated in a custom way. If the probability of detection was above the detection threshold, set in the INI file, its presence was marked red. Note a decent performance of the classifier in rats destroying their home cage.

At this stage, all 39 videos of the morning sessions and 36 videos of the evening sessions were stored; their number was reduced later due to the lack of some corresponding audio files at the final stage of processing.

### 3. M. Analysis of ultrasonic vocalizations

The analysis of audio files was conducted offline using DeepSqueak version 3.1.0, the program developed by Coffey et al., [16], on MATLAB (MathWorks, Inc., Natick, MA), version R2023a. The spectrograms were created using fast Fourier transform (FFT) with a window size = 0.0032 s, nfft = 0.0032 s and a 90% overlap. DeepSqueak uses a convolutional neural network for USVs detection. The 50- and 22-kHz USVs were detected separately.

The 50-kHz USVs were detected using multidetect function by its default detection network “Rat Detector YOLO R1”, frequency cutoff high: 95 kHz, frequency cutoff low: 35 kHz, score threshold: 0 (Precision = 0.97, Recall = 0.99, F1 = 0.98).

The 22-kHz USVs were detected using DeepSqueak’s default detection network “Long Rat Detector Yolo R1”, frequency cutoff high: 10 kHz, frequency cutoff low: 35 kHz, score threshold: 0.

The contour thresholds for all audio files were set to 0.215 for entropy and 0.825 for amplitude percentile. The detection files were processed using Post Hoc Denoiser to eliminate false positive results. Post Hoc Denoiser is a neural network capable of distinguishing USVs from common types of background noise and alarm reflections. The noise networks were trained in total, on 216 samples (3 audio files from the present experiment). For training purposes, the “Denoiser” samples were manually labeled as the calls or the noise (the validation accuracy = 81.48%, validation frequency = 10 iterations).

Following Post Hoc Denoising stage, all USV recordings were manually inspected and the inaccurate detections were corrected. DeepSqueak’s detections including inaccuracies (false positive and negative detections) were compared with the human detections on all USV calls using simple linear regression for all audio files.

The results of the regression analyses, shown in **Figure** 8, revealed significant similarities between human and DeepSqueak detections. The R^2^ values for 50-kHz and 22-kHz were 0.99 and 0.98, respectively (for both: P values < 0.0001).

Based on the acoustic call features, the 50-kHz calls were divided into the following types: short calls (mean call duration: <= 12 ms), flat calls (bandwidth: <= 6 kHz AND mean call duration: > 12 ms) and the frequency modulated calls (bandwidth: > 6 kHz AND mean call duration: > 12 ms) [25]. These calls were not subcategorized into steps, trills, etc. [26].

**Figure 8.**
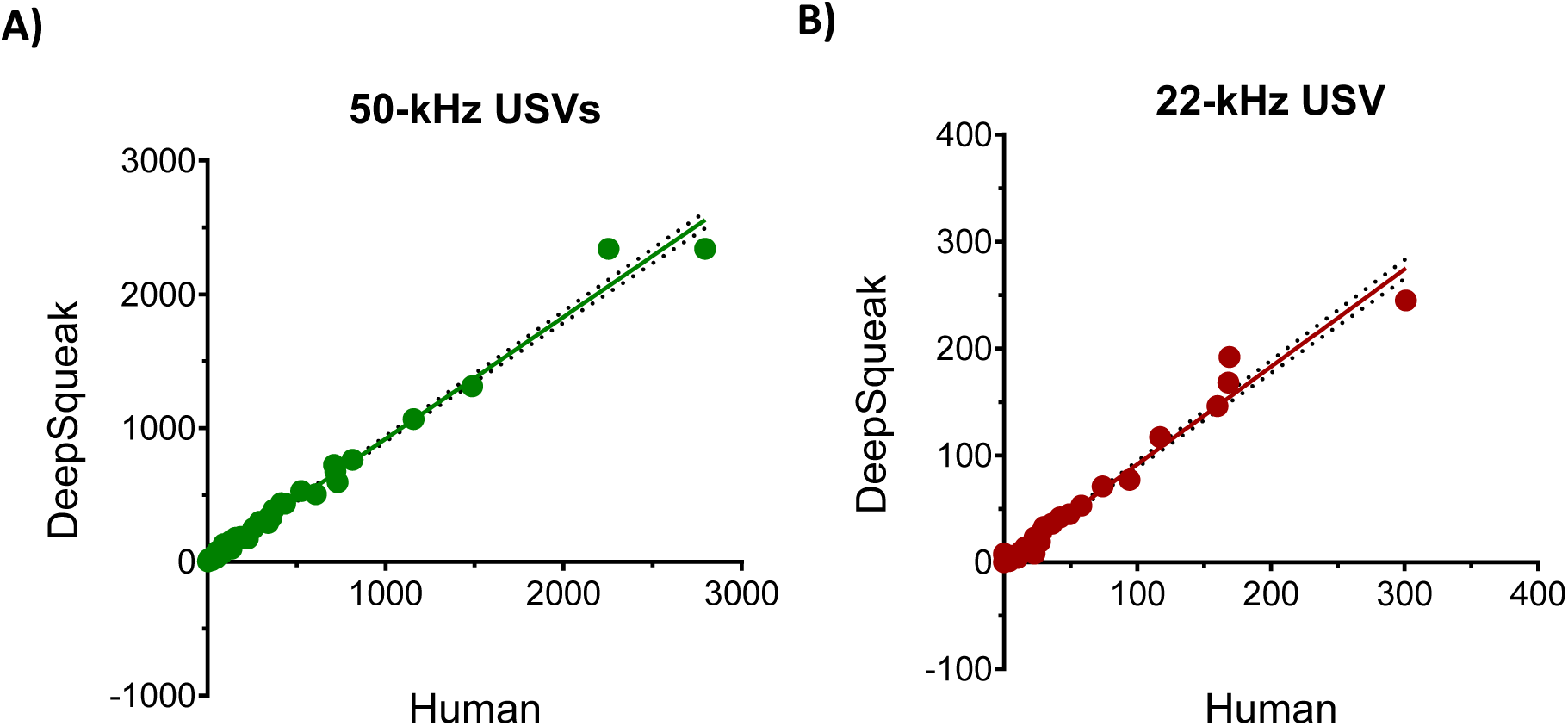
Regression analyses of the number of all ultrasonic calls identified by the experienced researcher (A.P.; “Human”) and by the DeepSqueak machine learning tool. Each point represents the number of USVs emitted by the pairs of rats in each of the 6 boxes recorded on 2 sessions (morning and evening) over 6 days of experiment (6 x 2 x 6 ∼ 72 audio files). As not all recordings were available, 34 “morning” and 37 “evening” audio files were analyzed, totaling to 71 measurements.

### 3. N. Analysis of associations between behavioral and ultrasonic categories

Beyond the analysis of rats’ activity at the two twilight periods, this work addressed also a more global question: if the ultrasonic vocalizations play a substantial role in rats’ social life, how strongly they are associated with a particular behavior and with other call type(s)?

To detect the behavior-USV call type, and the USV call type-USV call type associations, i.e., their co-occurrences, the analysis of social behavior videos (i.e., the SimBA’s “machine results” CSVs) and of USV categories (DeepSqueak’s output) had to be temporarily matched as precisely as possible.

As the videos were recorded at 12 frames per second (FPS), one frame of 1/12 seconds being represented as a single row in CSV files, constituted a single temporary unit, and could contain the behavior(s), USV call type(s), both, or none; see **Table** 2.

**Table 2.** explains the “frame”, i.e., the smallest time unit of 1/12 seconds. At a given time frame, a behavior, classified as absent (0) or present (1) was annotated, together with the presence or absence of a USV call type and their co-occurrence.

| Video frame number (1/12 seconds) | Was e.g., **fighting** behavior present? | Which of USV call types, if any, were present? | Were **alarm** calls present? | Were **flat** calls present? | Did **fighting** behavior and **flat** calls co-occur? |
| --- | --- | --- | --- | --- | --- |
| 1 | 0 | alarm<br>flat<br>alarm | 1 | 1 | 0 |
| 2 | 0 | alarm | 1 | 0 | 0 |
| 3 | 1 | flat | 0 | 1 | 1 |
| ... |  |  |  |  |  |

The construction of this dataset took several steps. First, the video and audio files stored at different computers, have had to be identified and their filenames have had to match. In this work, the main file with USV analysis (DeepSqueak output), with the columns including the date of observation, box name, exact begin and end times in relation to the beginning of the recording, the relevant *.wav file names and their paths, as well as its identified category (FLAT, SHORT, FREQ MODULATED and ALARM), contained > 18,000 rows. This file’s rows had to be first filled with the corresponding SimBA’s CSV video filenames.

### 3. O. Matching the video and audio filenames

In an ideal technical scenario – like with video recording on a smartphone – a video and a corresponding high quality (250 kHz) audio would be recorded simultaneously and accordingly enveloped. This was not the case of the present experimental setup because the video and audio recordings were done using different devices, stored at different computers/NAS stations, differed in their filenames and durations by more than several seconds.

Thus, a script^10^ analyzed the duration of videos stored in a custom folder, and based on FPS stored in video_info.csv SimBA’s file, it outputted frame numbers as well as the “Begin frame time (s)” and “End frame time (s)” columns in separate CSVs in the novel folder. The next script^11^ added USV types’ columns and filled them with zeros. Two helper scripts^12^,^13^ matched specific audio and video filenames.

### 3. P. Creating an index file listing all audio and video filenames

Following several manual adjustments, including deletion of duplicated rows, thanks to the script^14^, the index file was created and updated with the columns indicating the start and end of USV’s *.wav audio recordings as well as with their duration in seconds.

### 3. Q. Determining duration of audio files

The next script^15^ have read the updated index and SimBA’s video_info.csv files. Knowing the video’s FPS – needed for constructing N frames – and the precise date and time of the start and end of audio recordings, it outputted a novel set of CSVs files. These CSVs were of the number of rows equaling to the audio duration x FPS.

For instance, the CSV corresponding to the audio recording of duration of 1,200.005 seconds had 14,400 rows while a CSV corresponding to the audio recording of duration of 1,202.168 seconds (2 seconds longer) had 14,426 rows (26 more rows). The script, in addition created novel columns (Probability_ALARM, ALARM, Probability_FLAT, FLAT, Probability_FRQ MODUL., FRQ MODUL., Probability_SHORT, SHORT) and filled their cells with zeros. In short, thanks to this step we knew the precise beginning and the end of every audio file expressed as the frame numbers.

The next script^16^ updated audio CSVs in such a way that, if a specific call type was present at a given time of observation, the corresponding frame with that call type was updated with “1”. Thus, resulting CSVs had now “1” at rows where the call was detected; see **Table 2** above for an explanation. This was to create USV datasets based on DeepSqueak output because SimBA is intendent to be trained and to create the classifiers like “sniffing”, etc., rather than 1-frame indices of e.g., ALARM USV present. *Of note, we believe that the script could be adapted to broaden SimBA’s CSVs with any other (electrophysiological, telemetry, microdialysis, etc.,) numeric data that needed to be synchronized with the video recordings*.

### 3. R. Updating the index file with duration of individual videos

The next script^17^ filled out the video information columns in an updated index file. This index file was processed by the next script^18^, which compared the duration of audio and video files, and provided information whether the start and/or the end of the longer needed to be cut, to a) make the start of video and audio at the same exact moment and, b) of exactly same duration. This script saved the final index CSV file.

### 3. S. Adjusting the number of frames in audio CSVs and in video CSVs

Knowing the adjustments to be made, the CVSs representing audio data were saved thanks to the next script^19^. CSVs representing videos were optionally processed by the two scripts^20^,^21^ and then adjusted by script^22^. To make sure the length of video and audio was the same, as a **check point**, adjustments precision was visualized with the script^23^ that showed the longest mismatch of 1 second (12 frames), that for the media lasting for 1,200 seconds, appeared as an acceptable value.

### 3. T. Saving the final SimBA’s “machine_results” CSVs

The final set of CSVs with frames containing both SimBA’s detected behaviors and the novel USVs information, adjusted to the same duration, were stored in the temporary folder thanks to the next script^24^.

### 3. U. Adding human-hearable audio track to video files of the same start time and duration

Since we also aimed to merge the videos with human-hearable audios, the videos’ start and/or end were cut if necessary, with the script^25^ and stored in a novel temporary folder.

Then, thanks to ffmpeg’s custom function:

’lowpass=65000,highpass=20000,asetrate=25000, atempo=2,atempo=2,atempo=2,atempo=1.25,volume=3’,

the audio *.wav files were converted to human-hearable *.aac files with the script^26^ and stored in a separate folder. The *.aac files’ start and/or end were then cut, if necessary, with the script^27^.

As a **check point**, thanks to the next script^28^, the shortened videos received labels indicating behavioral categories as well as USV types superimposed on videos.

The final videos, merged with corresponding audios (**Figure** 9), were created with the next script^29^ that stored audible videos in a novel folder.

**Figure 9.**
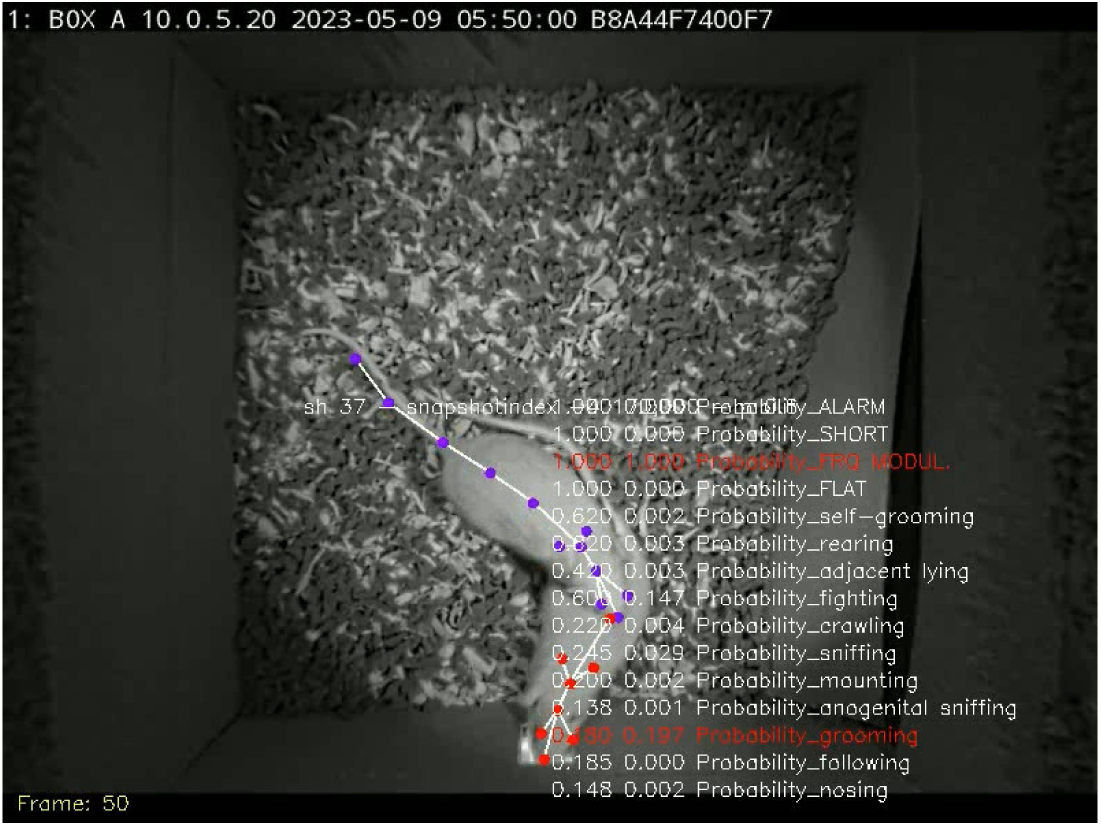
A frame from the final video containing human-hearable audio track showing grooming behavior and FM USV call at frame #50

### 3. V. Analysis of the number and duration of behaviors and ultrasonic calls over 1-minute epochs and days of observation

The scripts used in the following sections were created in python 3.12 environment.

The database of behaviors, USV types, and their co-occurrences (the Excel file) presented in 20, (1-minute) epochs, was produced by the script^30^. While SimBA offers a way to “stich” the episodes into the “bouts”, we were interested in recalculating them as the time, because SimBA’s INI file was defined to count every episode of nosing, sniffing etc., as one frame (see **Table** 2 above). The episodes were converted manually into duration (seconds of “nosing”, “sniffing”, etc., per 1-minute epoch, i.e., were divided by FPS = 12).

We decided to use 1-minute “epochs” to simplify data presentation, as the whole observation lasted for 20 minutes. For the “morning” and “evening” sessions, the minutes 1-10 represented the dark, and the light phases, respectively. If a behavior was observed continuously during the whole 1-minute epoch, it could take at the maximum 720 frames (60 seconds x 12). In practice, we observed that rats could display only “adjacent lying” behavior continuously for one minute.

Shortly, the script have read CSVs represented in the index file, assigned behaviors and USV call types into the relevant 1-minute epochs and saved the results in another Excel file. The next script^31^ processed that file and saved it as a novel Excel file, assigning the days of experiment to the actual dates, as the video and audio data were not available for all 6 days (**Figure** 10 and **Figure** 11).

**Figure 10.**
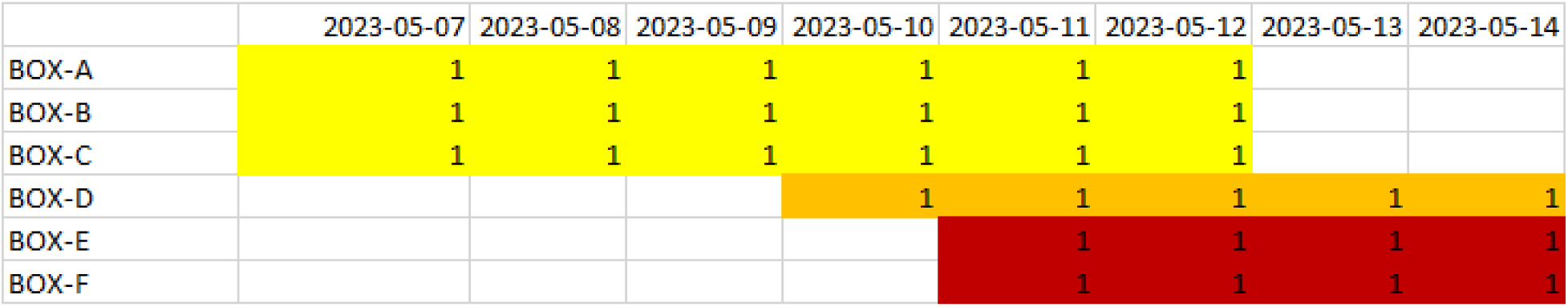
The morning sessions “calendar” showing data availability (“1”) for different dates and boxes.

**Figure 11.**
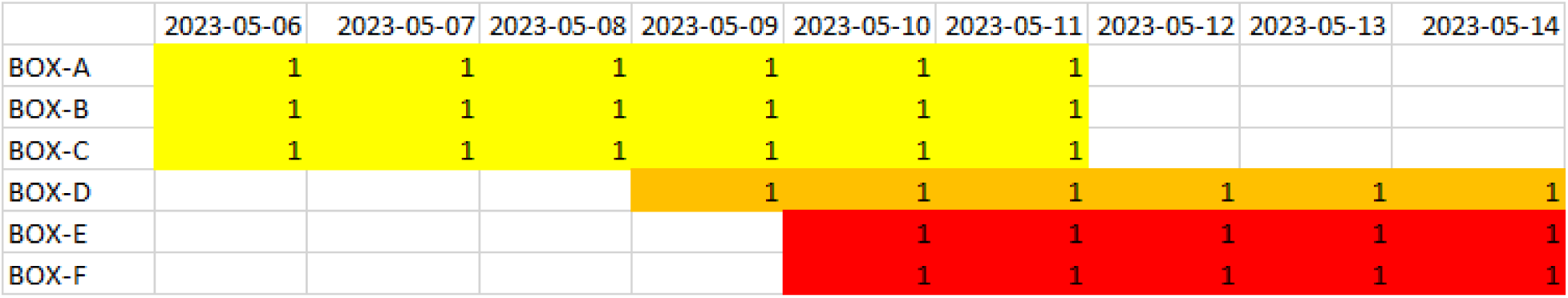
The evening sessions “calendar” showing data availability (“1”) for different dates and boxes.

Script^32^ have read the novel Excel file and outputted individual small CSVs with behaviors and USV category times in seconds per epoch and per day of experiment.

### 3. W. Plotting 3-D graphs

The scripts^33^ have read small CSVs from the previous step, and plotted individual 3-D graphs, examples of which is shown below:

**Figure 12.**
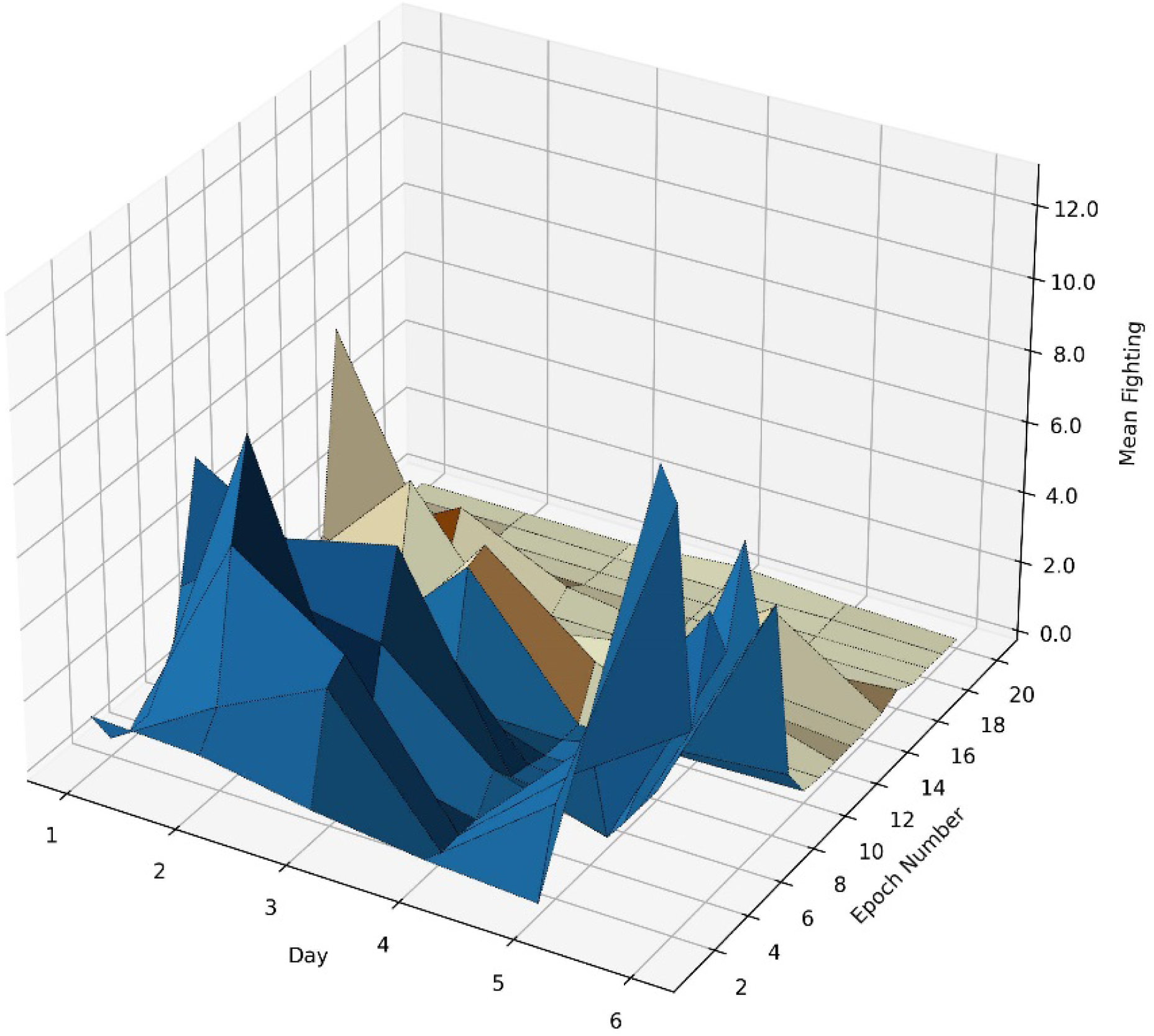
A 3-D plot showing the mean time (s) of fighting behavior in the morning sessions at different days, in 1-minute epochs. Blue and yellowish parts represent the dark and the light phase, respectively; minute 11 is time 06:00.

**Figure 13.**
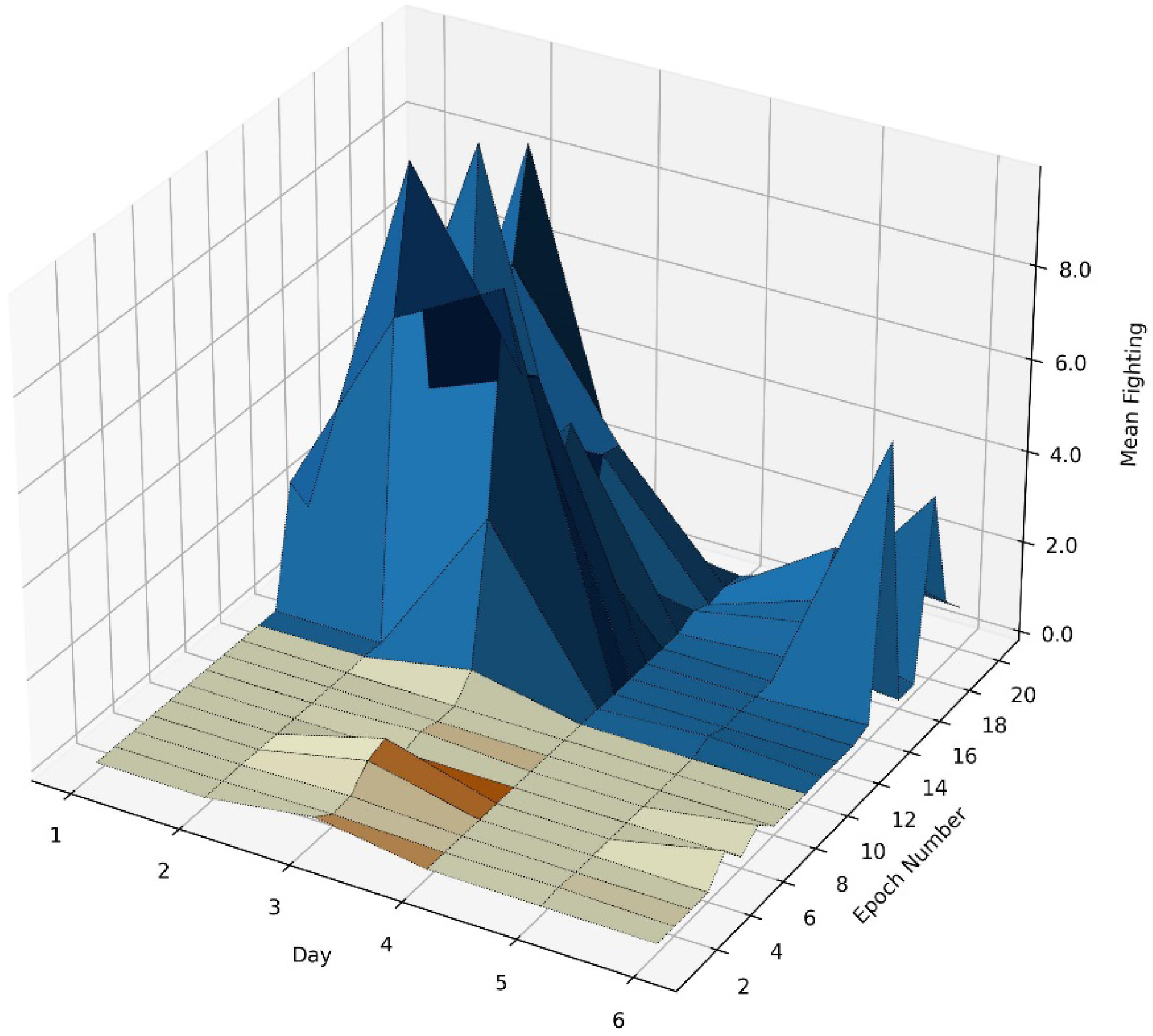
A 3-D plot showing the mean time (s) of fighting behavior in the evening sessions at different days, in 1-minute epochs. Yellowish and blue parts represent the light and the dark phase, respectively; minute 11 is time 18:00.

### 3. X. Statistics

We hypothesized that at the very moment the lights in animal housing boxes were turned off, the intensity of social behaviors would increase. Behavioral and ultrasonic data could be analyzed in various ways, as they were collected every 1/12 seconds for 20 minutes in the morning and in the evening sessions, over 4-6 days. However, the paucity of data expressed in frames precluded their analysis with repeated measures ANOVA. Moreover, the detailed analysis of each frame would include 10 min x 720 = 7,200 frames before and 7,200 frames after light change.

When inspecting 3-D figures plotted in 1-minute intervals, it was apparent that there is a clear difference between and after light change. To further simplify statistical analyses and graphs, for every box (A-F), data recorded before and after light change were averaged. Similarly, since not all data existed for all days, for every box, the data of the early phase (days 1-3) and late phase (days 4-6) were averaged. While not ideal, this step allowed for construction of 2 datasets represented as epochs (before-after light change) and of 2 datasets represented as early phase (days 1-3) and late phase (days 4-6) of the experiment, for each box.

Thus, data expressed as the duration (s) of 11 behavioral and of 4 ultrasonic categories were first checked for normality using Anderson-Darling, D’Agostino & Pearson, Shapiro-Wilk, and Kolmogorov-Smirnov tests. As the most of them failed to show normal distribution, data were rank-transformed [27], and subjected to two-way, repeated measures ANOVA with epoch (before-after light change), and early/late phase of experiment, for each box. Boxes (A-F) served as subjects. To determine whether the change in lighting conditions affected intensity of social behavior and of USV categories, and whether the phase of experiment played a role, 15 x 2 = 30 repeated measures analyses of variance were performed using IBM SPSS ver. 29. The alpha level was set at 0.05.

### 3. Y. Analysis of the associations between behaviors and USV call types, and between types of USVs

We hypothesized that certain behaviors (like fighting) are associated with some USV call types (like aggressive 22-kHz calls). Scripts^34^ saved another Excel file, with data aggregated for the days 1-3 and 4-6 and for the epochs 1-10 and 11-20. Scripts^35^ have read this Excel file and created the contact sheets with all co-occurrences. These plots represented the sums of a behavior and USV type or, two types of USVs. For instance, if in a given 1-minute epoch there was 1 “short” USV call and 4 “frequency modulated” USV calls, the sum of their co-occurrences was 5.

### 3. Z. Quality system

At the time of performing this study, Department of Behavioral Neuroscience and Drug Development has been holding EQIPD [28]; see https://go-eqipd.org/ certificate no. PL-INS-DBNDD-11-2021-2 (Nov 12th, 2021 – Nov 30th, 2024); see https://paasp.net/new-eqipd-certified-research-unit/ and https://go-eqipd.org/about-eqipd/eqipd-certified-research-groups/. Thus, this study followed EQIPD guidelines: https://eqipd-toolbox.paasp.net/wiki/EQIPD_Quality_System for instance, it was carefully designed in advance, and was documented in detail at every stage of progress.

## 4. Results

### 4. A. General activity of rats as measured by continuous monitoring of head entries into the water supply

Daily pattern of rats’ general activity showed increased number of head entries at the dark phase as compared with the light phase. Two way ANOVA with data collapsed for dark/light phase demonstrated differences between phases: F(1,5314)=236; P<0.001, **Figure** 14.

**Figure 14.**
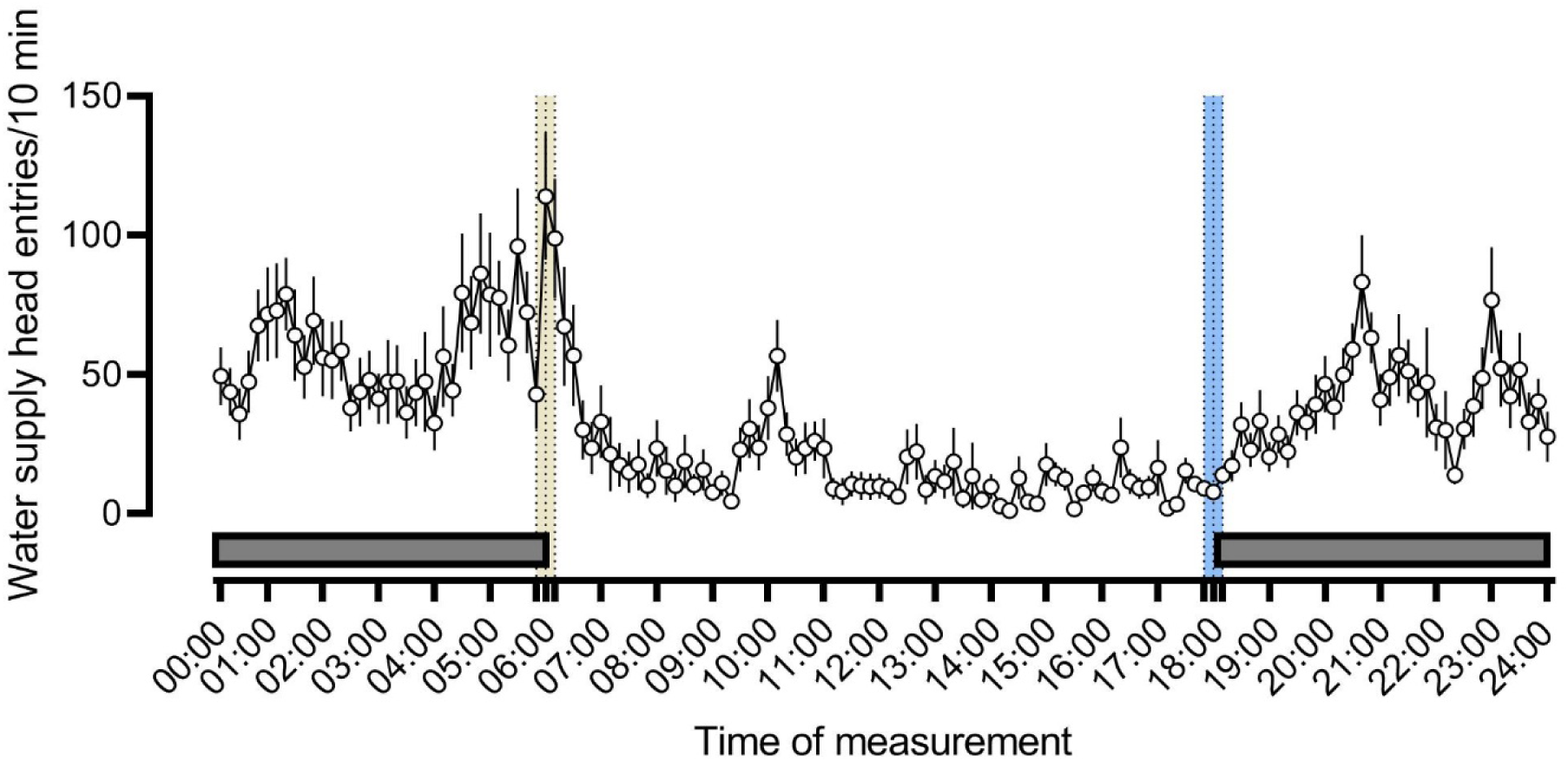
Daily pattern of water supply head entries of pairs of rats in 10-minutes epochs. Each point represents mean +/-SEM of 3-6 experimental boxes monitored over 5-7 days of experiment. This measure is a proxy of animals’ activity, likely but not necessarily reflecting water drinking, as we observed rats placing their heads into the lickometer while sniffing, or just lying near to the water tube. The black horizontal bars represent the dark phase (06:00-18:00). The yellow bar shows the 20-min “morning” sessions, beginning at 05:50 and ending at 06:10; the central dotted line indicating 06:00 hour, the end of darkness. The blue bar shows the 20-min “evening” sessions, beginning at 17:50 and ending at 18:10; the central dotted line indicating 18:00 hour, the beginning of darkness. Pairs of rats were monitored between days of 2023-05-05 and 2023-05-15.

A separate ANOVA carried out on four 10-minutes epochs (05:50-06:00, 06:00-06:10, 17:50-18:00, and 18:00-18:10) also demonstrated differences among groups: F(3,146)=12.83; P<0.001, with data of the time 06:00-06:10 being different from the 05:50-06:00 time point, **Figure** 15.

**Figure 15.**
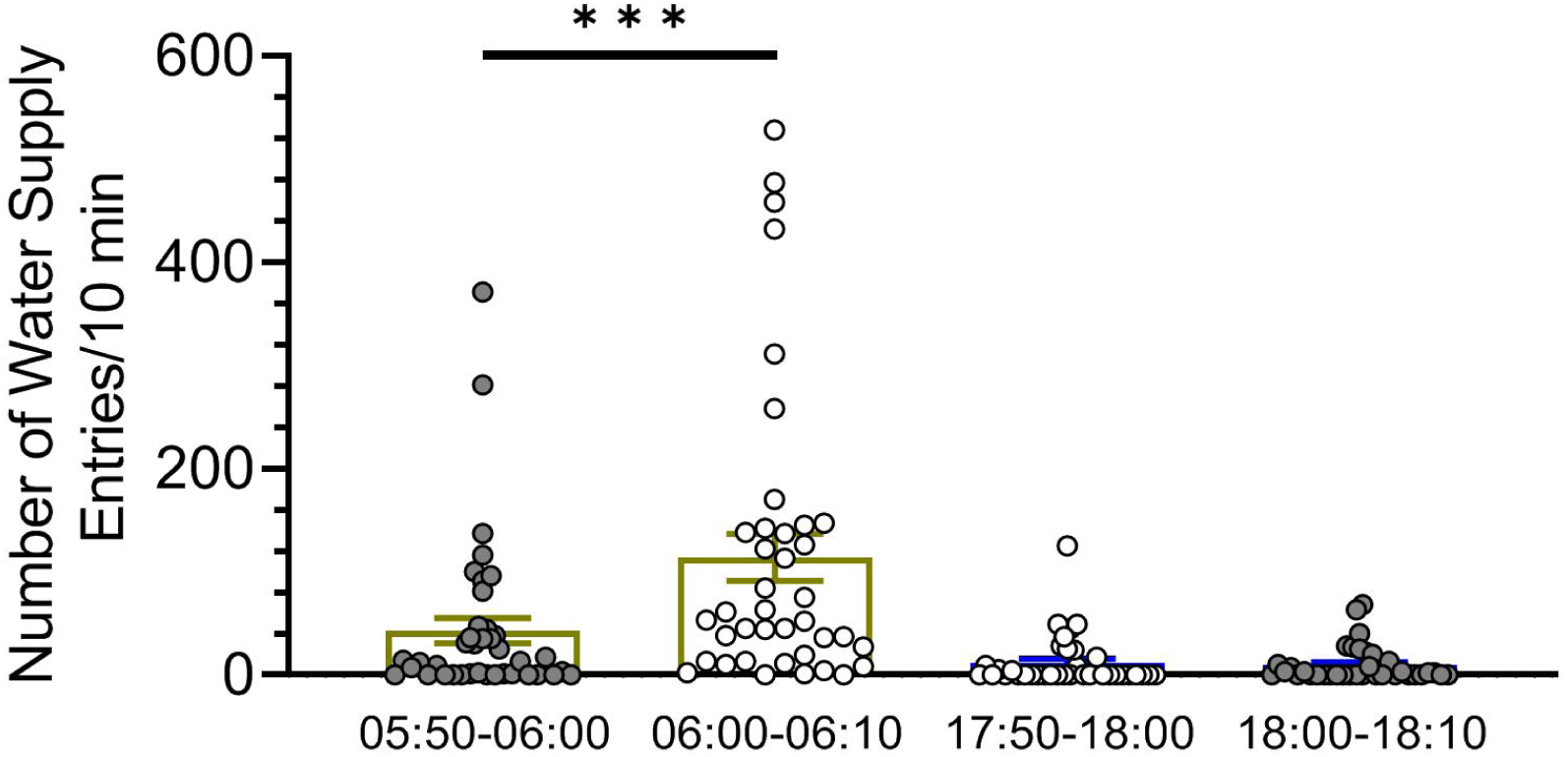
Water supply head entries, a raw measure of rat’s activity shows rapid increase at the time the lights were turned on at 06:00 but no change at the time the lights were turned off. Symbols: *** P<0.001, 06:00-06:10 vs. 05:50-06:00, Tukey HSD.

### 4. B. Duration of rats’ social behaviors and ultrasonic calls at the beginning of the light phase (morning sessions)

In the morning sessions, we observed light-on induced increases in anogenital sniffing, adjacent lying and rearing behavior.

In addition, rats displayed less of grooming and rearing as well as less of alarm, frequency modulated and short calls at days 4-6 than at days 1-3.

**Table 3.** Statistical details of data presented on Figure 16.

| Behavioral or USV category | Effect of the Day of experiment<br>[Partial Eta Squared] | Effect of the lights turned ON<br>[Partial Eta Squared] |
| --- | --- | --- |
| Nosing | $F < 1$ | $F(1,5)=1.99$; NS |
| Following | $F < 1$ | $F(1,5)=4.38$; NS |
| Grooming | $F(1,5)=16.81$; $P=0.009$ [0.771] | $F < 1$ |
| Fighting | $F(1,5)=3.57$; NS | $F(1,5)=3.29$; NS |
| Anogenital sniffing | $F < 1$ | $F(1,5)=8.98$; $P=0.030$ [0.643] |
| Mounting | $F < 1$ | $F(1,5)=1.08$; NS |
| Sniffing | $F(1,5)=3.54$; NS | $F(1,5)=2.57$; NS |
| Crawling | $F < 1$ | $F(1,5)=2.77$; NS |
| Adjacent lying | $F(1,5)=5.47$; NS | $F(1,5)=9.32$; $P=0.028$ [0.651] |
| Rearing | $F(1,5)=12.36$; $P=0.017$ [0.712] | $F(1,5)=13.39$; $P=0.015$ [0.728] |
| Self-grooming | $F < 1$ | $F(1,5)=1.17$; NS |
| ALARM calls | $F(1,5)=7.19$; $P=0.044$ [0.590] | $F < 1$ |
| FLAT calls | $F(1,5)=6.42$; NS | $F < 1$ |
| FREQUENCY MODULATED (FM) calls | $F(1,5)=7.105$; $P=0.045$ [0.587] | $F < 1$ |
| SHORT calls | $F(1,5)=15.24$; $P=0.011$ [0.753] | $F < 1$ |

**Figure 16.**
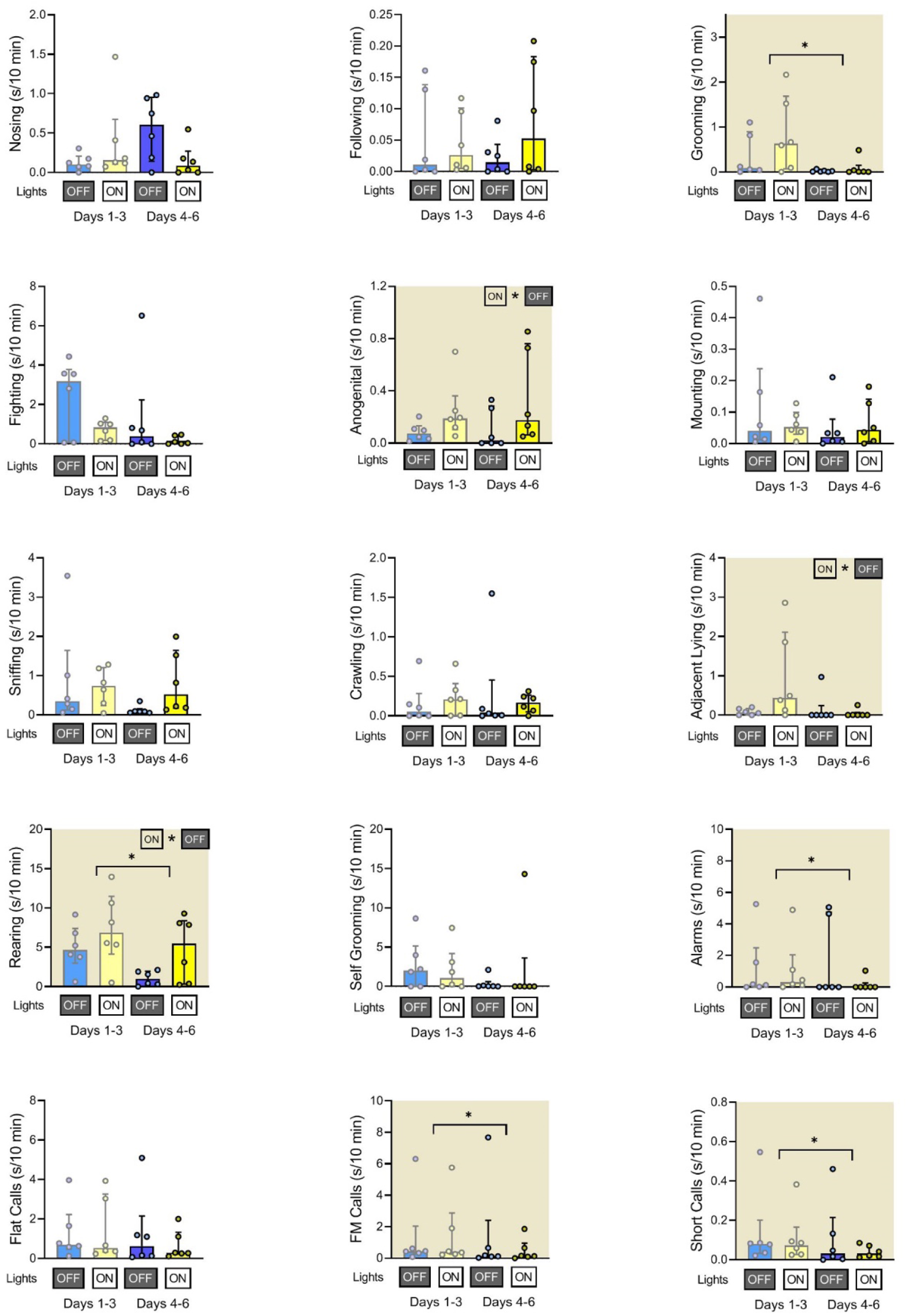
Duration of rats’ social behaviors and ultrasonic calls at the morning sessions. The blue bars indicate the end (05:50-06:00; lights off) of the dark phase and yellow bars indicate the beginning (06:00-06:10; lights on) of the light phase. The early (days 1-3) and late (days 4-6) of the experiment are shown separately. Plots with shaded background indicate that the early/late days factor (horizontal line) and/or that the lights on/off factor (ON * OFF) was significant (P<0.05). Data are presented as median +/-interquartile range.

### 4. C. Duration of rats’ social behaviors and ultrasonic calls at the end of the light phase (evening sessions)

In the evening sessions, we observed light-off induced increases in fighting, mounting, crawling, rearing (but not in anogenital sniffing and adjacent lying as was observed in the morning sessions) as well as of the alarm, flat and short calls (not observed at the morning sessions).

In addition, rats displayed less of flat (not observed in the morning sessions) as well as of frequency modulated and short calls (like in the morning sessions) at days 4-6 than at days 1-3.

**Table 4.** Statistical details of data presented on Figure 17.

| Behavioral or USV category | Effect of the Day of experiment<br>[Partial Eta Squared] | Effect of the lights turned OFF<br>[Partial Eta Squared] |
| --- | --- | --- |
| Nosing | F(1,5)=4.57; NS | F(1,5)=2.75; NS |
| Following | F(1,5)=5.05; NS | F(1,5)=1.95; NS |
| Grooming | F=1 | F=1 |
| Fighting | F(1,5)=2.89; NS | F(1,5)=18.37; P=0.008 [0.786] |
| Anogenital sniffing | F(1,5)=3.38; NS | F < 1 |
| Mounting | F(1,5)=2.25; NS | F(1,5)=12.71; P=0.016 [0.718] |
| Sniffing | F < 1 | F < 1 |
| Crawling | F < 1 | F(1,5)=9.43; P=0.028 [0.654] |
| Adjacent lying | F < 1 | F < 1 |
| Rearing | F(1,5)=3.42; NS | F(1,5)=7.99; P=0.037 [0.615] |
| Self grooming | F < 1 | F < 1 |
| ALARM calls | F < 1 | F(1,5)=23.48; P=0.005 [0.824] |
| FLAT calls | F(1,5)=11.75; P=0.019 [0.702] | F(1,5)=36.48; P=0.002 [0.879] |
| FREQUENCY MODULATED (FM) calls | F(1,5)=8.43; P=0.034 [0.628] | F(1,5)=3.71; NS |
| SHORT calls | F(1,5)=9.59; P=0.027 [0.657] | F(1,5)=18.42; P=0.008 [0.787] |

**Figure 17.**
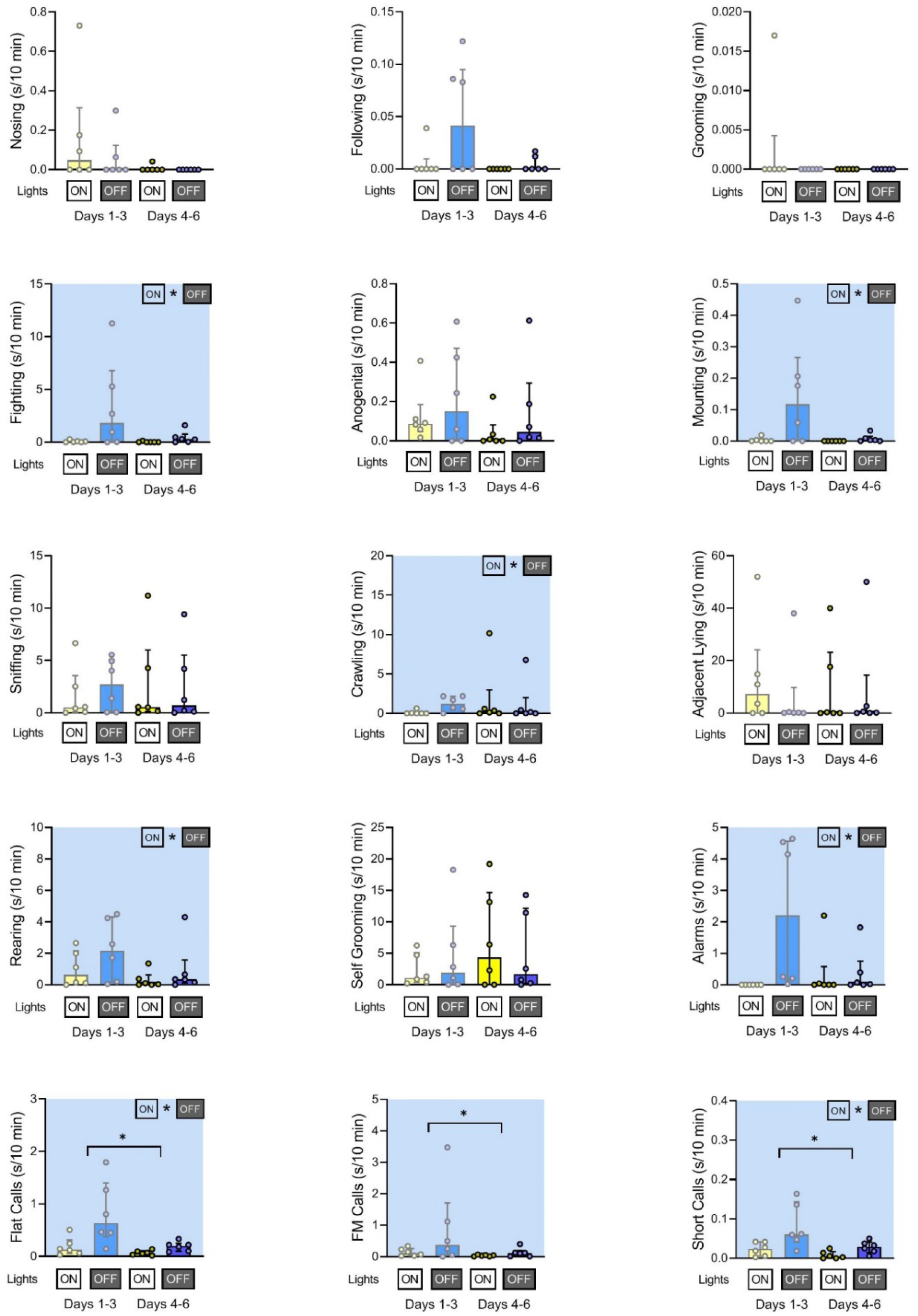
Duration of rats’ social behaviors and ultrasonic calls at the evening sessions. The yellow bars indicate the end (17:50-18:00; lights on) of the light phase and blue bars indicate the beginning (18:00-18:10; lights off) of the dark phase. The early (days 1-3) and late (days 4-6) of the experiment are shown separately. Plots with shaded background indicate that the early/late days factor (horizontal line) and/or that the lights on/off factor (ON * OFF) was significant (P<0.05). Data are presented as median +/-interquartile range.

### 4. D. Associations between ultrasonic calls and social behaviors at the beginning of the light phase (morning sessions) of pairs of rats presented at the twilight (dark/light phase) over early (days 1-3) and late (days 4-6) phase of experiment

The analyses of associations between ultrasonic calls and social behaviors could be summarized as follows. During the morning sessions, alarms were associated with the flat and frequency modulated calls. The alarm, flat, frequency modulated and short calls were associated with the rearing behavior, mostly at the beginning of the light phase.

**Figure 18.**
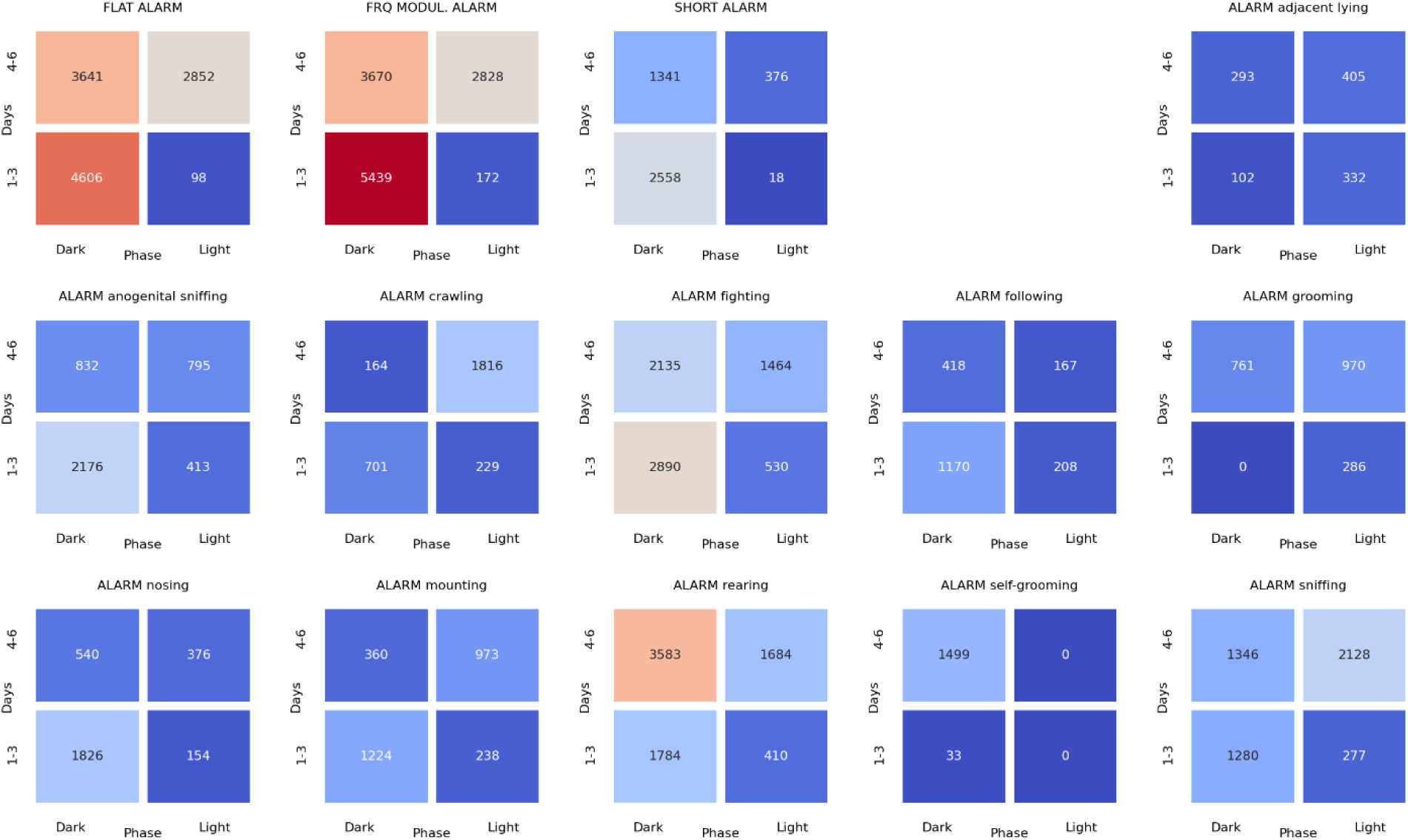
The associations of the **alarm** ultrasonic calls with 3 other USV call types as well as with 11 behaviors recorded over early (1-3) and late (4-6) days of the **morning** sessions, analyzed separately for the beginning and the end of the twilight. As shown, these calls were associated with the flat and frequency modulated calls at the dark phase, especially at the early days. The alarm calls were also associated with the rearing behavior during the dark phase of the late days.

**Figure 19.**
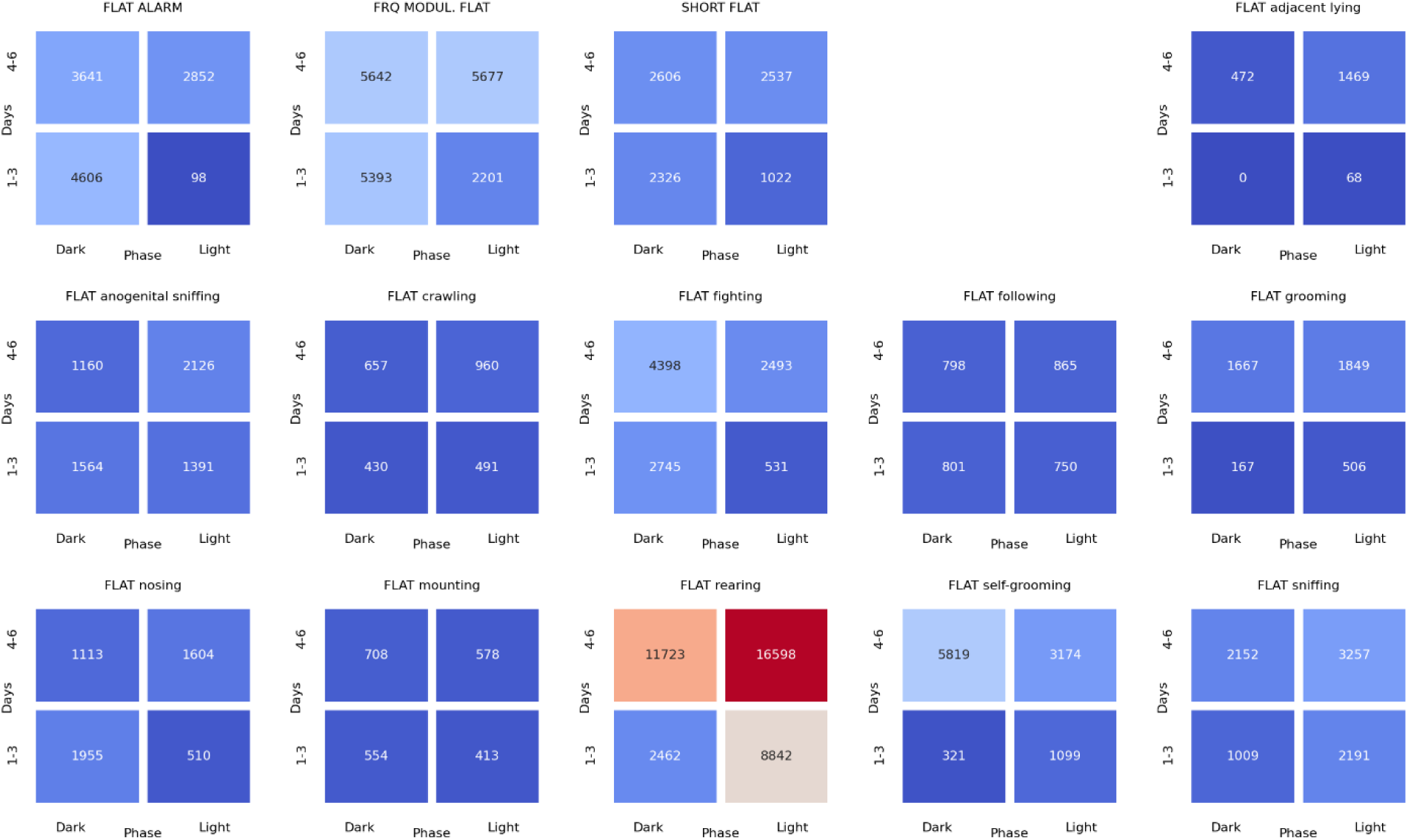
The associations of the **flat** ultrasonic calls with 3 other USV call types as well as with 11 behaviors recorded over early (1-3) and late (4-6) days of the **morning** experiment, analyzed separately for the beginning and the end of the twilight. As shown, these calls were associated with the rearing behavior during the dark and light phases of the late days.

**Figure 20.**
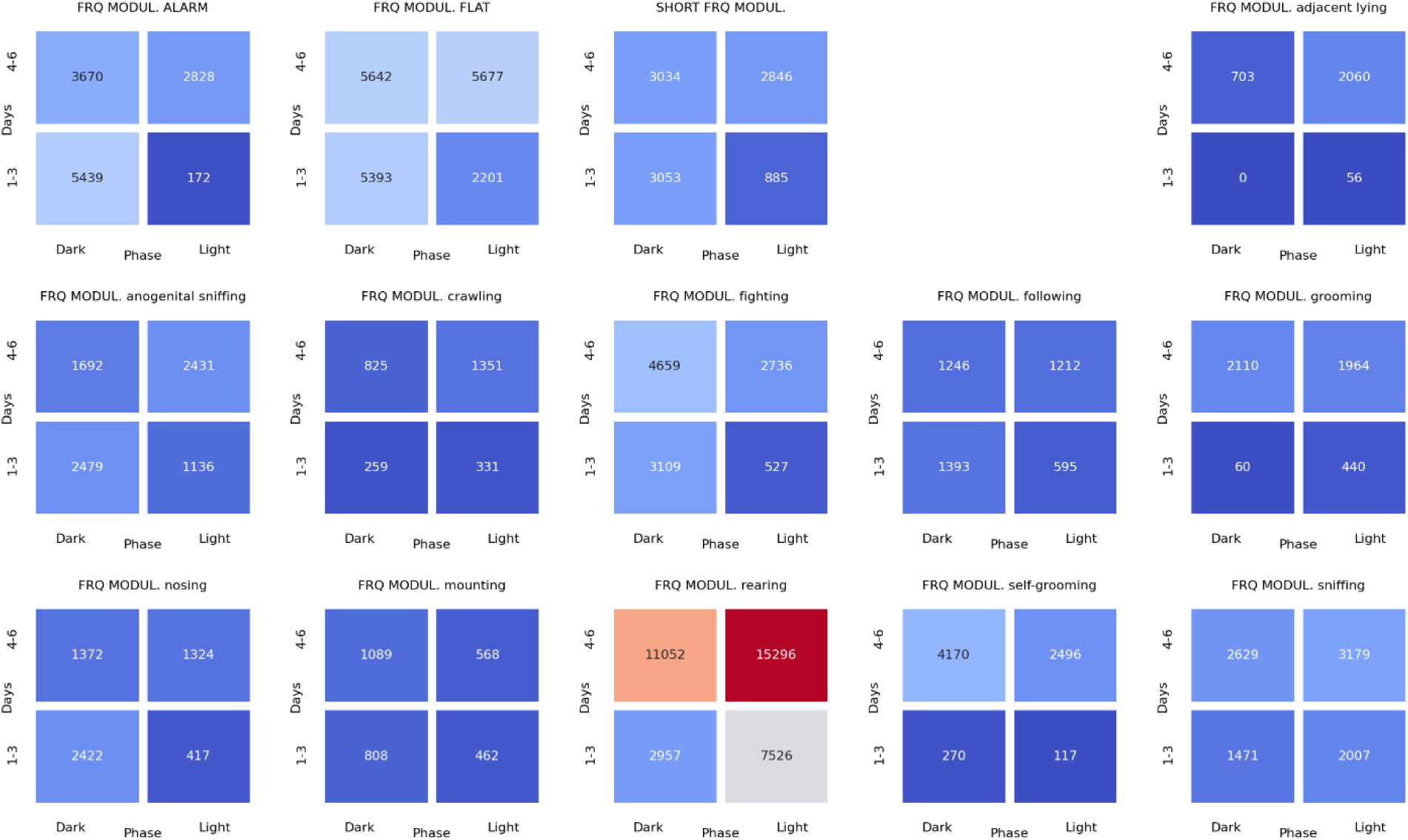
The associations of the **frequency modulated** ultrasonic calls with 3 other USV call types as well as with 11 behaviors recorded over early (1-3) and late (4-6) days of the **morning** experiment, analyzed separately for the beginning and the end of the twilight. As shown, these calls were associated with the rearing behavior during the dark and light phases of the late days.

**Figure 21.**
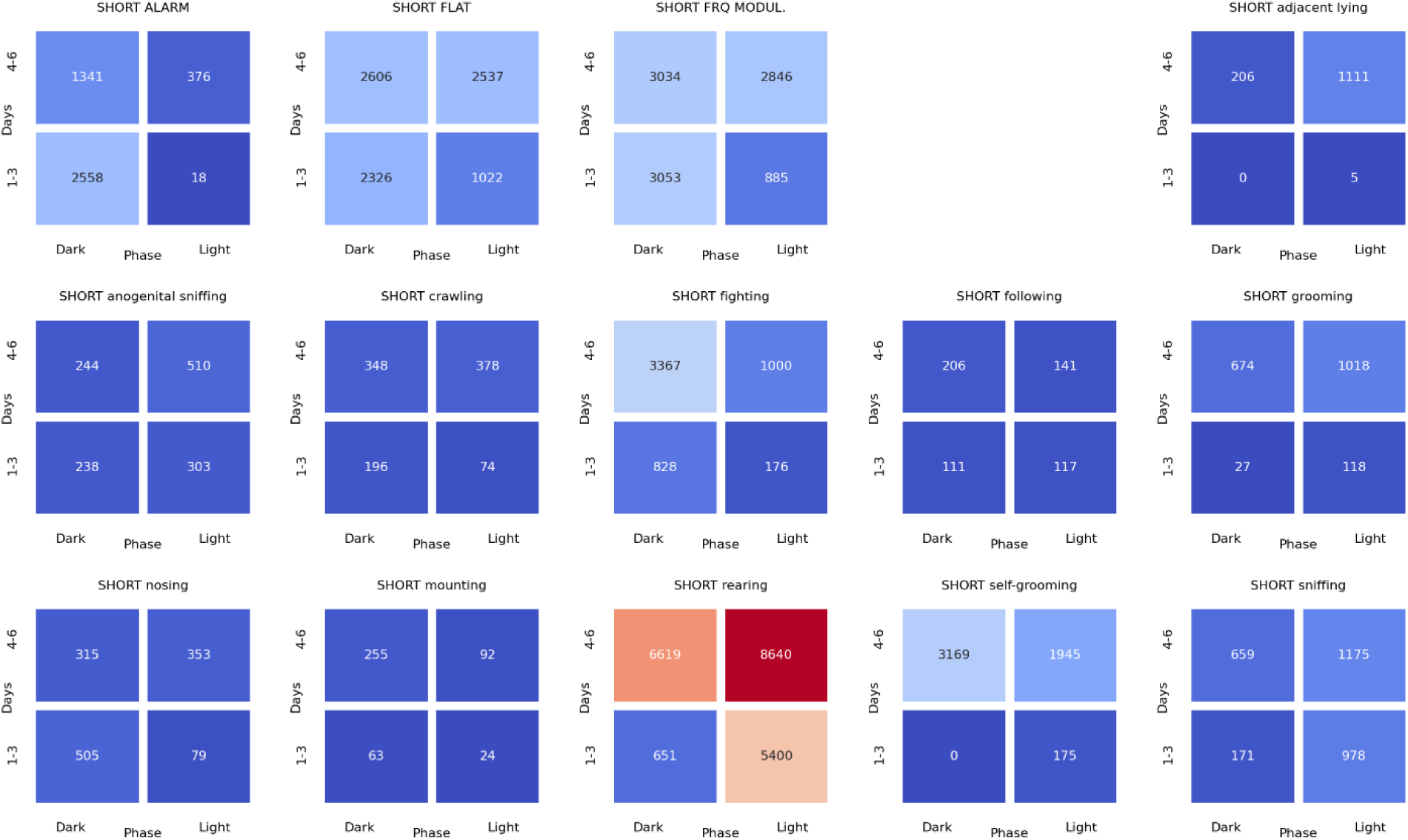
The associations of the **short** ultrasonic calls with 3 other USV call types as well as with 11 behaviors recorded over early (1-3) and late (4-6) days of the **morning** experiment, analyzed separately for the beginning and the end of the twilight. As shown, these calls were associated with the rearing behavior.

### 4. E. Associations between ultrasonic calls and social behaviors at the beginning of the dark phase (evening sessions) of pairs of rats presented at the twilight (light/dark phase) over early (days 1-3) and late (days 4-6) phase of experiment

The analyses of the associations between ultrasonic calls and social behaviors could be summarized as follows. During the evening sessions, the alarm, flat, and frequency modulated calls were associated with other types of calls, but also with sniffing, crawling, fighting, and rearing. Of note, the flat and short calls were associated with the adjacent lying.

**Figure 22.**
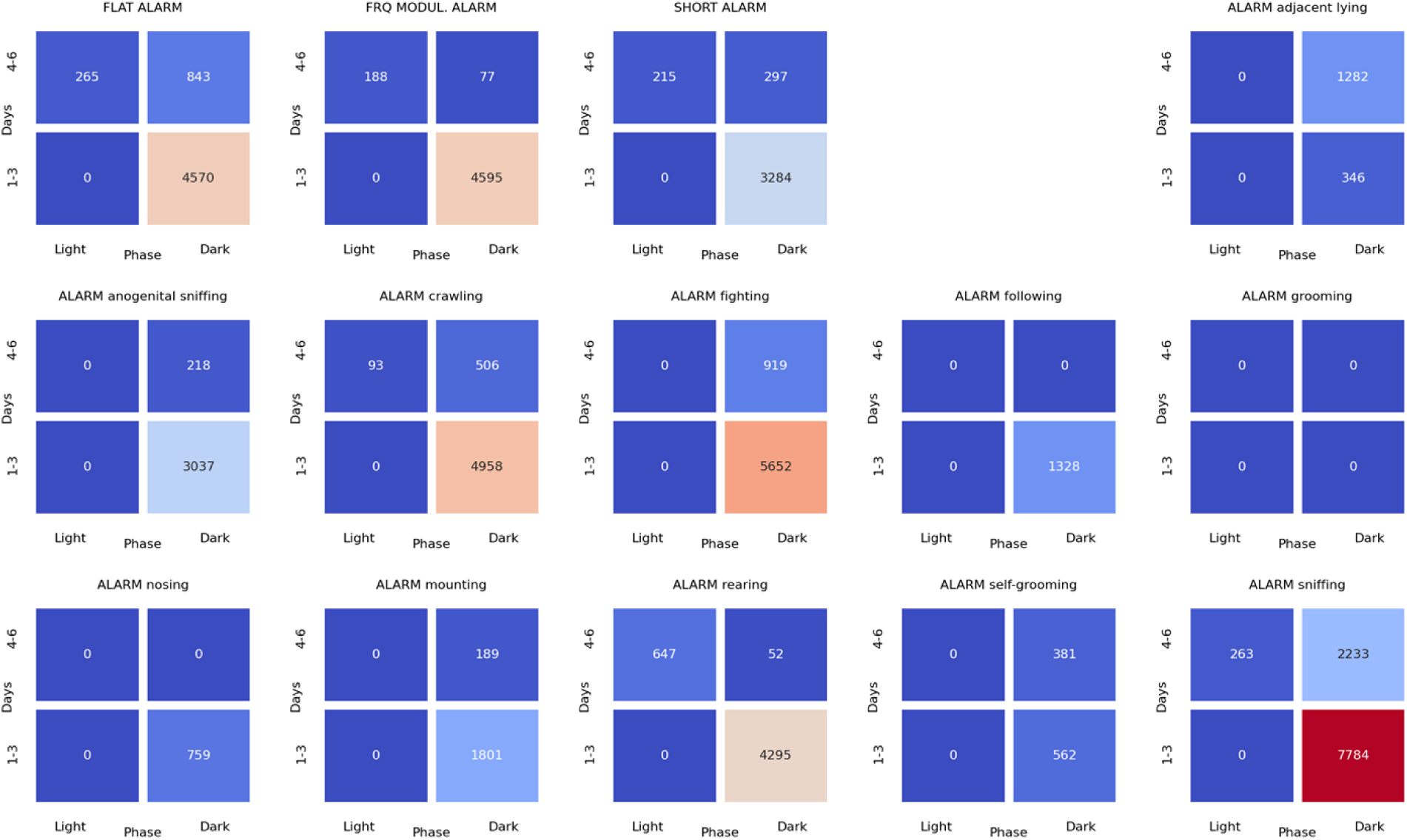
The associations of the **alarm** ultrasonic calls with 3 other USV call types as well as with 11 behaviors recorded over early (1-3) and late (4-6) days of the **evening** sessions, analyzed separately for the beginning and the end of the twilight. As shown, these calls were associated with the flat and frequency modulated calls at the dark phase, especially at the early days. The alarm calls were also associated with the crawling, fighting, sniffing, and rearing behaviors during the dark phase of the early days.

**Figure 23.**
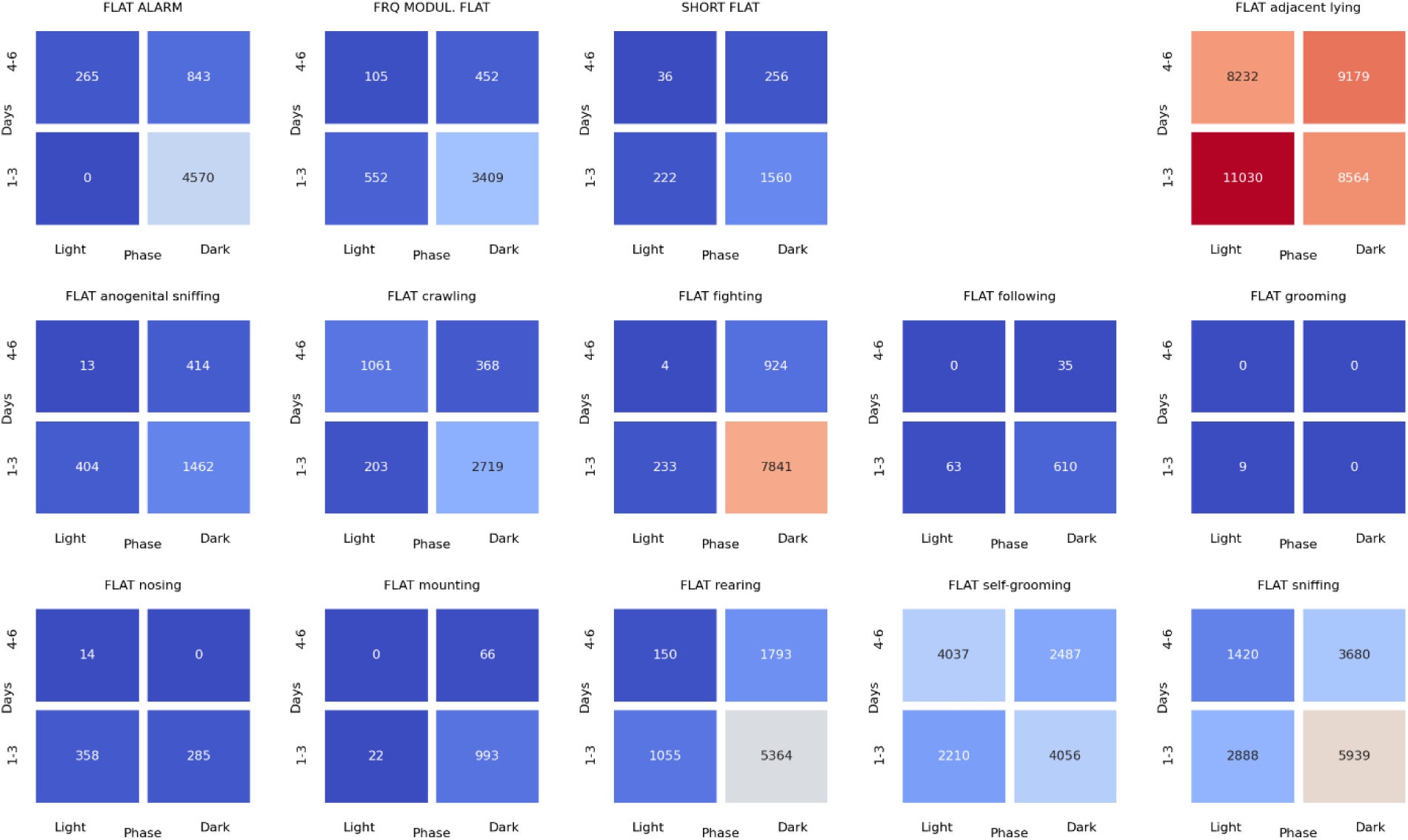
The associations of the **flat** ultrasonic calls with 3 other USV call types as well as with 11 behaviors recorded over early (1-3) and late (4-6) days of the **evening** experiment, analyzed separately for the beginning and the end of the twilight. These calls were associated mostly with the adjacent lying behavior. We also observed that the flat calls were associated with fighting, rearing and sniffing behaviors, at the beginning of the dark phase during early days.

**Figure 24.**
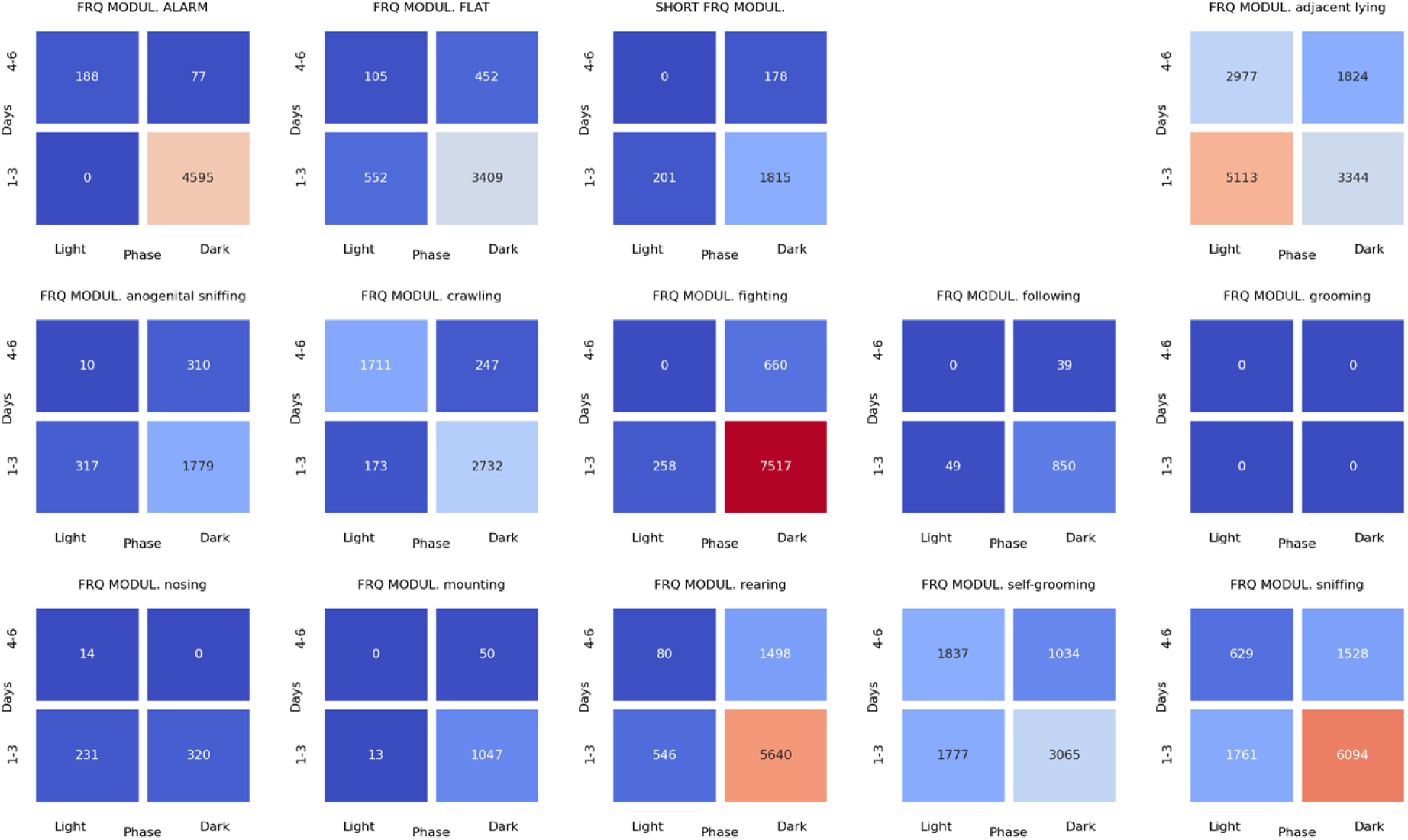
The associations of **frequency modulated** ultrasonic calls with 3 other USV call types as well as with 11 behaviors recorded over early (1-3) and late (4-6) days of the **evening** experiment, analyzed separately for the beginning and the end of the twilight. These calls were associated with the fighting, sniffing, and rearing behaviors at early days during the beginning of the dark phase. The FM calls were also associated with the alarm calls and adjacent lying behavior at the early days of experiment.

**Figure 25.**
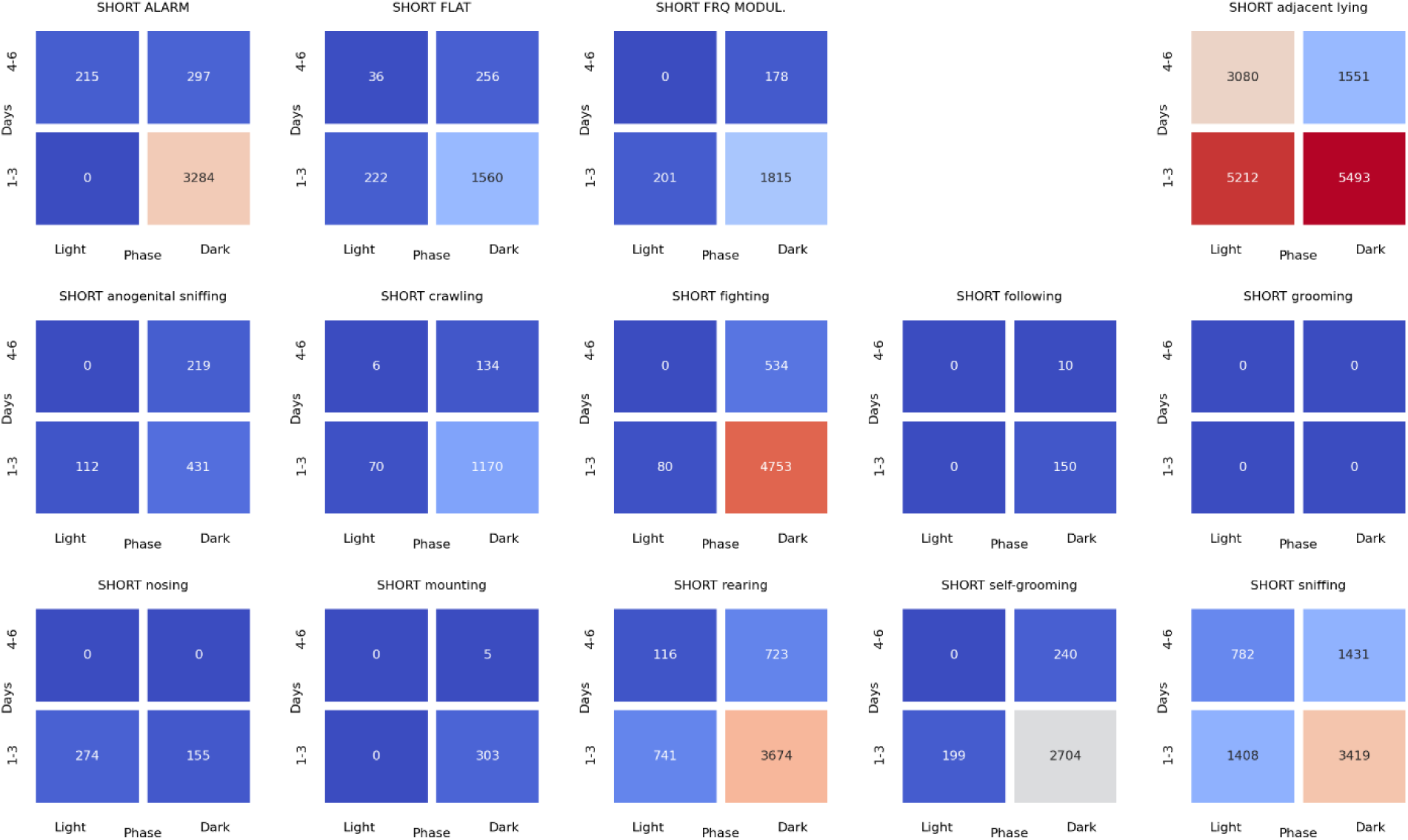
The associations of the **short** ultrasonic calls with 3 other USV call types as well as with 11 behaviors recorded over early (1-3) and late (4-6) days of the **evening** experiment, analyzed separately for the beginning and the end of the twilight. These calls were associated mostly with adjacent lying behavior. Short calls were associated with the fighting, **rearing and** sniffing behaviors, at the beginning of the dark phase at the early days of experiment.

## 5. Discussion

Humans display diurnal fluctuations in the positive affect. It is unclear, when exactly they happen, as there are reports showing either the strongest peak in positive affect occurring in the morning or in the night [29, 30]. In the wild and laboratory rats, the daily rhythm of activity, measured as the percentage of rats active outside the nest, [2] and as the positive affect, exemplified by the number of “hedonic” 50-kHz ultrasonic calls, appears to be maximal during the active lights-off period [5], showing, in addition, the early and late activity cycle peaks. Burgdorf et al., [5] further suggested that the peaks may translate to an ability to coordinate pro-social play behavior right after the start of the wake cycle and to complete pro-social behavior right before the end of the wake cycle.

The aim of the present work was to verify a common observation that the laboratory rats intensify their activity at the moment the lights in the animal facility were turned off. As the phenomenon of darkness-induced prosocial activity has an apparent heuristic value, we were curious whether it could be investigated using the top modern digital techniques. Specifically, we examined the nature of darkness-induced activity; in that, whether it affects the social life and if so, which behavioral categories were altered. In addition, we investigated whether darkness-induced activity is associated with the ultrasonic vocalizations; the “happy” (50-kHz) and/or “alarm” (22-kHz) calls.

Present study confirms that at the dark phase, the animals were more active as measured by water supply head entries (**Figure** 14). A more detailed analysis, however, revealed that, contrary to our predictions, right after the lights were turned off, the animals did not start to poke into the lickometers. Conversely, they significantly increased the water supply head entries right after the lights were turned on (**Figure** 15). While this phenomenon could be explained by light-induced agitation, it suggests that this simple way of monitoring animals’ activity will unlikely reveal darkness-induced activity. The discrepancy between present and Stryjek et al. [2], results could be due to the way the data were collected. While the lickometeres monitored every nose poke, and then data were aggregated into 10-minutes epochs, Stryjek et al., [2] observed every fifth photo from those taken once a minute by a video camera.

The analysis of evening sessions confirms that at the beginning of darkness, the rats increased their activity [2], but the increase did not concern social behavior categories in the expected way, see **Figure** 17 and summary **Table** 5. In the evening, the **social investigatory** behaviors including nosing, following, anogenital sniffing and sniffing were not affected. The darkness also did not affect **amicable** grooming [22], and self-grooming [24]. In contrast, with the lights-off, the animals started to display increased fighting, mounting and crawling behavior, the two latter being associated with and supporting **aggression** [14, 15, 21–23]. These behaviors were concomitant with several forms of USVs, including alarm, flat, and short calls. Of note, the **frequency-modulated** “happy” or “hedonic” calls were not affected by the darkness. Together with increased fighting, increased 22-kHz alarm calls strongly suggest that when the rats wake up, they start to display aggressive behavior. As expected, we observed no increase in **adjacent lying**, which, while amicable, is a way the rats sleep or rest together [19, 20]; something that, logically, would not increase at the beginning of the active phase of the light-dark period. Darkness-induced activity was also due to increased rearing behavior, which represents a form of environmental **exploration**, and with the flat and short calls, associated with investigative behavior and general arousal [31].

**Table 5.** The summary of findings regarding the effects of rapid change in light intensity on duration of social behaviors and ultrasonic calls. Two far right columns show how the social life adapts (decreases) at days 4-6 compared to days 1-3 of the experiment. Yellow columns indicate the morning and the blue columns indicate the evening sessions, respectively.

| Behavioral or USV category | Lights-on effect | Lights-off effect | Late days, morning | Late days, evening |
| --- | --- | --- | --- | --- |
| Nosing |  |  |  |  |
| Following |  |  |  |  |
| Grooming |  |  | ↓ |  |
| Fighting |  | ↑ |  |  |
| Anogenital sniffing | ↑ |  |  |  |
| Mounting |  | ↑ |  |  |
| Sniffing |  |  |  |  |
| Crawling |  | ↑ |  |  |
| Adjacent lying | ↑ |  |  |  |
| Rearing | ↑ | ↑ | ↓ |  |
| Self-grooming |  |  |  |  |
| ALARM calls |  | ↑ | ↓ |  |
| FLAT calls |  | ↑ |  | ↓ |
| FREQUENCY MODULATED (FM) calls |  |  | ↓ | ↓ |
| SHORT calls |  | ↑ | ↓ | ↓ |

Analysis of behaviors recorded at the evening sessions revealed also that animals adapt to the novel home cages. As stated above, we tested pairs of rats continuously, for 6 days in a semi-natural conditions, which were novel to the animals. The establishment of social hierarchy takes time [32, 33], and one would predict that some behaviors, especially related to aggression, could display lower intensity at the later (4-6) days than early (1-3) days of experiment. However, at the evening sessions we failed to show such adaptation for any behavioral category, though we noted that the 50-kHz USVs of were reduced at the late days. This did not concern 22-kHz alarm calls, associated with aggressive behaviors.

The morning sessions served as internal control conditions. When the lights were turned on, and the animals were supposedly going to sleep, one would expect a decrease in social activities. Indeed, we noted an increase in the resting activity (adjacent lying; [19, 20]), consistent with decreased activity at high light level, see **Figure** 16 and summary **Table** 5. However, as with the evening sessions, the exploratory activity (**rearing**) increased, likely due to the agitation induced by a rapid change in light intensity. Lights-on did not affect any category of USVs. Analysis of the morning sessions also revealed adaptation to the novel home cages. At the later (4-6) days we observed less of **grooming** and **rearing** behaviors and of ultrasonic calls than at early (1-3) days of experiment. Of note, none of other behaviors’ duration was decreased over the course of experiment in the morning sessions.

The second issue examined addressed a more global question: if the ultrasonic vocalizations indeed play a substantial role in rats’ social life, how strongly they are associated with a particular behavior? For instance, are 22-kHz “alarm” calls associated with the fighting behavior, as it is commonly [34, 35] but not universally [36] accepted? Both during the morning (**Figure** 18) and the evening (**Figure** 22) sessions, the **alarm** calls were indeed associated with the fighting, crawling, mounting and sniffing behavior, see summary **Figure** 26. This again supports the notion that the alarm calls communicate aggressive behavior [34, 35]. The 22-kHz calls were also associated with other calls’ types and with the rearing behavior, suggesting a complex pattern of ultrasonic communication.

The **frequency modulated** “happy” or “hedonic” calls were associated with investigatory sniffing and rearing behaviors as well as with adjacent lying and fighting. This pattern suggests that they co-occur with the amicable, but also with aggressive behaviors. A similar pattern of associations was noted for the **flat and short** calls, that were, in addition, associated with the adjacent lying, see summary **Figure** 26. As non-frequency modulated flat and short calls were postulated not to bear the positive affect [37–41], their presence at times the rats were resting together is not surprising.

**Figure 26.**
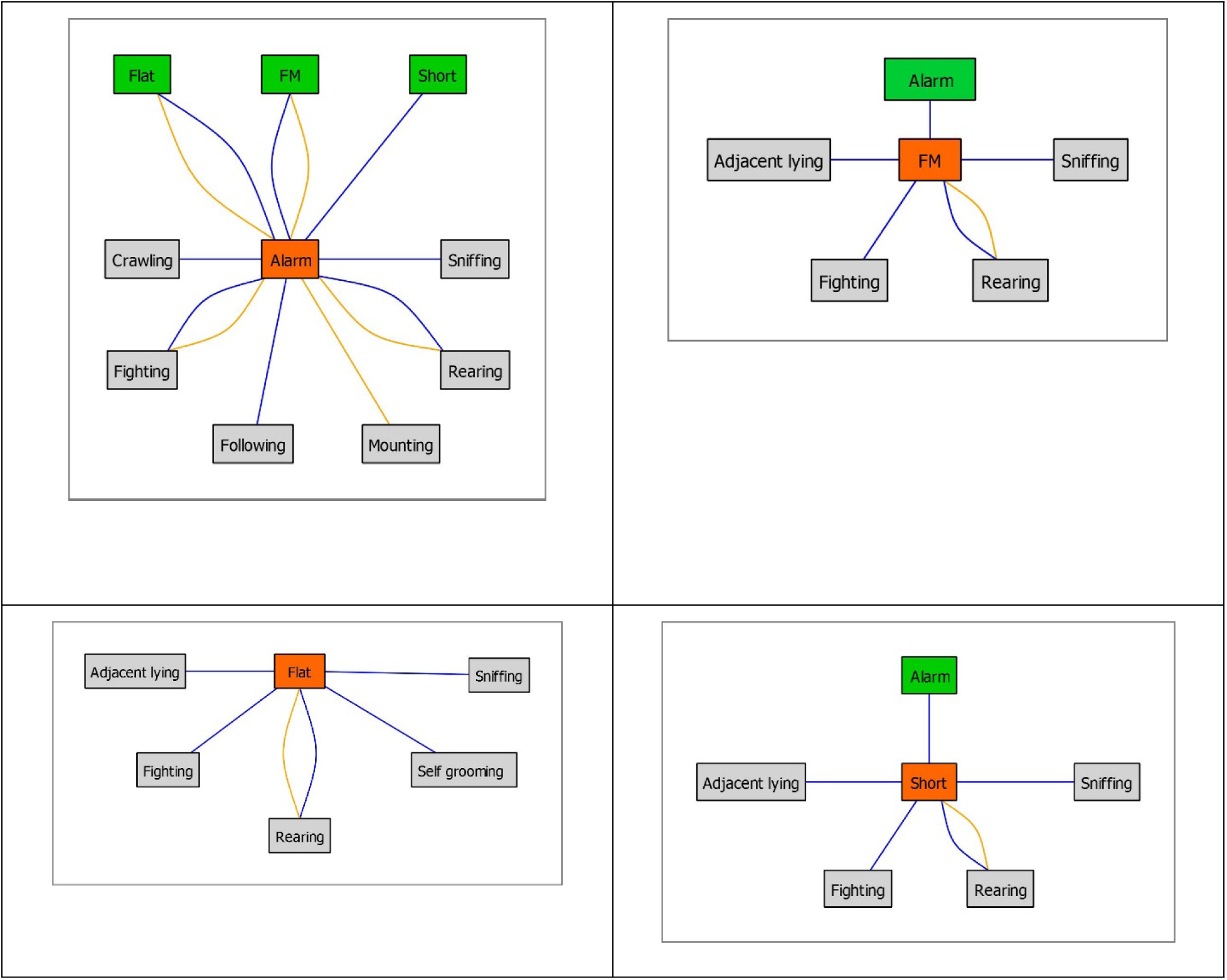
Summary of the associations between ultrasonic calls and social behavior categories presented on Figures 18-25. Blue and yellow lines indicate the evening and morning sessions’ associations, respectively. The green boxes show other ultrasonic categories associated with the main (red) ultrasonic call type.

Altogether, one may hypothesize that the lights-on stimulus agitates the animals inducing rearing behavior and ultrasonic vocalizations. This is consistent with the increase in general activity, represented by water supply head entries (**Figure** 15). However, present results do not support the hypothesis that the lights-off stimulates pro-social behavior and hedonic calls. This might be due to several reasons. The discrepancy between present ultrasonic data and Burgdorf et al., [5] results could be due to technical reasons, because while we calculated the total time of USVs, Burgdorf et al., [5] measured their number. Alternatively, the animals could have been displaying aggressive behaviors and alarm calls because they were continuously establishing the social hierarchy [33, 42]. However, we did not observe significant adaptation (decrease) of the fighting behaviors and alarm calls at the evening sessions over the days of experiment, though it is debatable how long it takes for the rats to establish the dominance status [43, 44]. As we observed a global decrease of social behavior over the course of experiment, chances were that observing the rats for longer than 6 days would decrease even further their social activities. One also cannot exclude that the aggressive behavior is indeed rewarding to the aggressor [45], as not only the alarm, but also flat and short (though not frequency modulated) 50-kHz calls, were increased. Unfortunately, the present experimental setup disallowed for distinguishing which animal in a pair was calling. This apparent limitation could be overcome using more sophisticated than ours, ultrasonic “camera” devices [46, 47].

Present work provides several novel utilities and observations. Experimental setup involving custom build inexpensive boxes allowed for simultaneous recording of video, audio, and general activity. Using several python’s scripts, we were able to as precisely as possible, merge the ultrasonic information with the videos, and we believe that a slight modification of the workflow would allow to synchronize other types of data. The videos supplemented with the human-hearable audio track allow for a direct observation of rats’ behavior. The DeepSqueak machine learning tool [16], together with other free tools including DeepLabCut (for reviews see [17, 48]) as well as the SimBA open-source package [14, 15] allowed for rapid and objective analysis of ultrasonic and behavioral data. The DeepSqueak analyses were similar to the results generated by an experienced experimenter, and much faster as compared to the human workflow. This laboratory communicated recently a remarkable similarity of rat social behavior annotations made by an experienced researcher’s and DeepLabCut [9].

In conclusion, we present a study with an easy to replicate step-by-step workflow allowing objective measurement of rat social behavior and ultrasonic vocalizations. Regardless of the nature of darkness-induced activity, we hypothesize that it could be compromised in apathy-like states associated depressive-like phenotype, allowing for studying anti-apathy medications. As spontaneous apathy or anergia awaits for a valid animal model to be studied in semi-natural conditions, these hypotheses are currently tested in our laboratory.

## 6. Supplementary files

Supplementary files including python’s scripts are available at http://gofile.me/5TcdS/l9XirUEnz.

## 7. STAR Methods

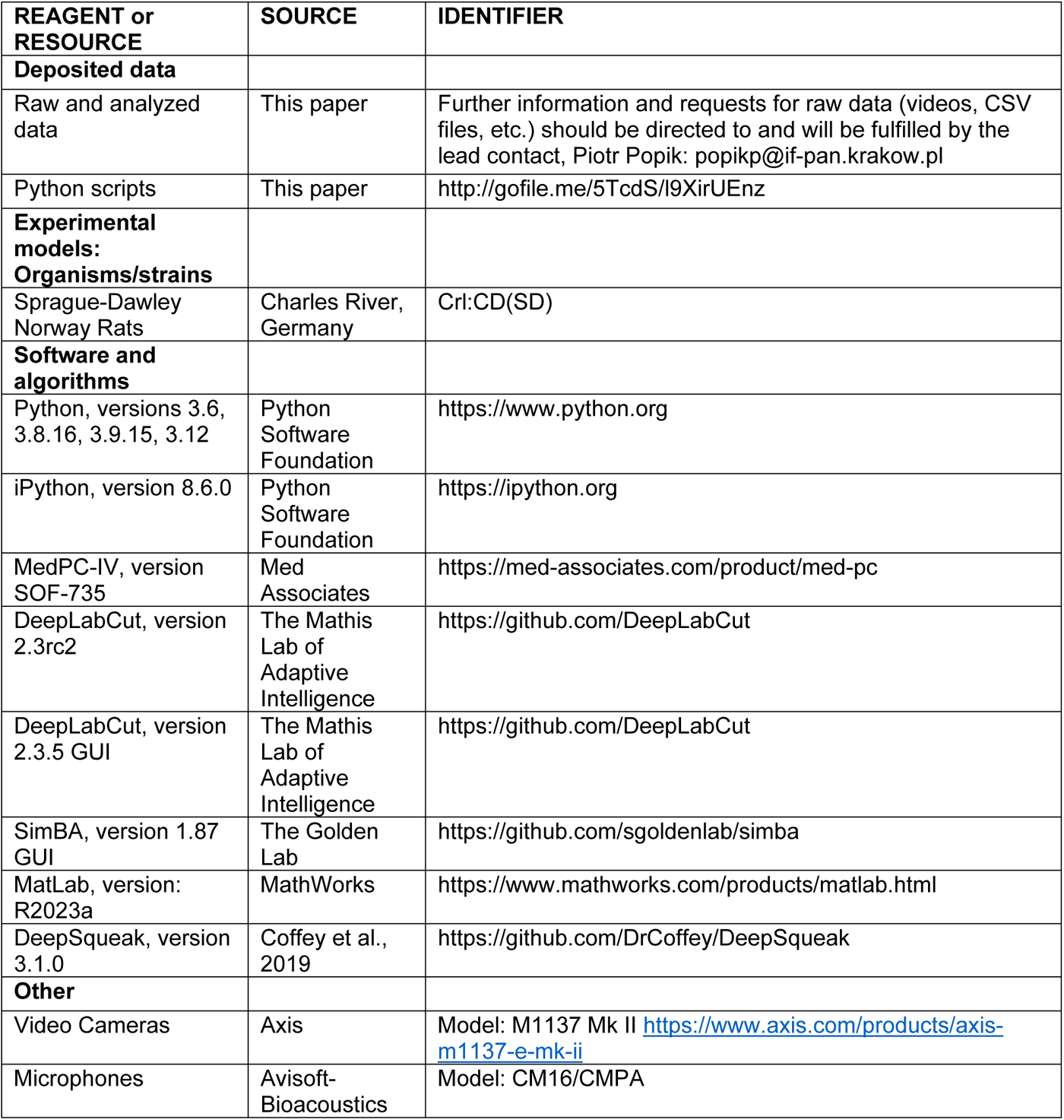
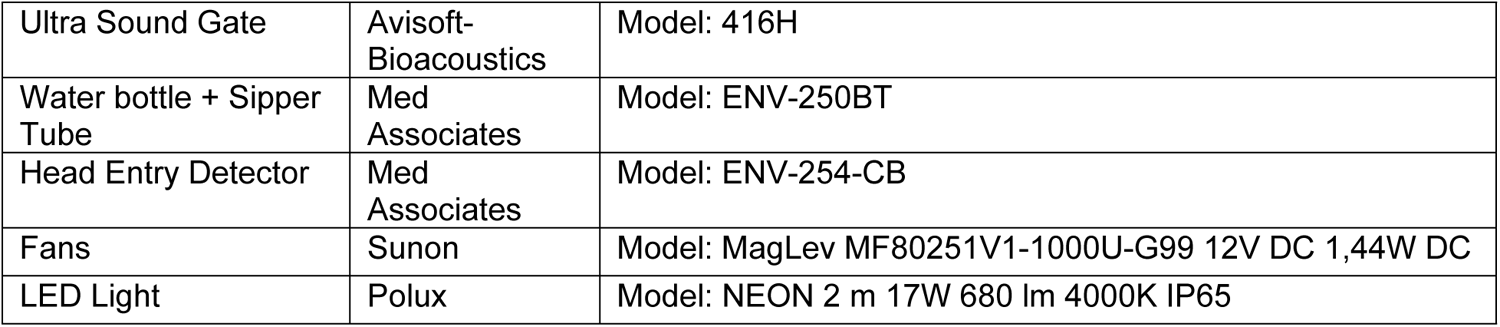

## Acknowledgements

This work could not be done without availability of open source DeepLabCut https://github.com/DeepLabCut marker-less pose-estimation and Simple Behavioral Analysis (SimBA) https://github.com/sgoldenlab/simba toolkits. For these, the authors deeply thank the DeepLabCut Team: Alexander Mathis and Mackenzie Mathis, Tanmay Nath and Jessy Lauer, the DeepLabCut Community as well as the Golden Laboratory: Simon Nilsson and Sam A. Golden, respectively. We thank Kevin R. Coffey for DeepSqueak, a deep learning based system for detection and analysis of ultrasonic vocalizations. Cephares https://www.cephares.pl/ project is acknowledged for providing the Nvidia DGX A100 computer station.

## Author contributions

PP designed research; AN, NMR, AP and JG performed research; PP and EC analyzed data; PP wrote the paper.

## Data Availability Statement

Data that support the findings of this study are available at http://gofile.me/5TcdS/l9XirUEnz.

## Footnotes

1 01-popik_et_al_20240629.py

2 02-popik_et_al_20240629.py

3 03-popik_et_al_20240629.py

4 04-popik_et_al_20240629.py

5 05-popik_et_al_20240629.py

6 06-popik_et_al_20240629.py

7 07-popik_et_al_20240629.py

8 08-popik_et_al_20240629.py

9 09-popik_et_al_20240629.py

10 10-popik_et_al_20240629.py

11 11-popik_et_al_20240629.py

12 12-popik_et_al_20240629.py

13 13-popik_et_al_20240629.py

14 14-popik_et_al_20240629.py

15 15-popik_et_al_20240629.py

16 16-popik_et_al_20240629.py

17 17-popik_et_al_20240629.py

18 18-popik_et_al_20240629.py

19 19-popik_et_al_20240629.py

20 20-popik_et_al_20240629.py

21 21-popik_et_al_20240629.py

22 22-popik_et_al_20240629.py

23 23-popik_et_al_20240629.py

24 24-popik_et_al_20240629.py

25 25-popik_et_al_20240629.py

26 26-popik_et_al_20240629.py

27 27-popik_et_al_20240629.py

28 28-popik_et_al_20240629.py

29 29-popik_et_al_20240629.py

30 30-popik_et_al_20240629.py

31 31-popik_et_al_20240629.py

32 32-popik_et_al_20240629.py

33 33E-popik_et_al_20240629.py or 33M-popik_et_al_20240629.py

34 34E-popik_et_al_20240629.py or 34M-popik_et_al_20240629.py

35 35E-popik_et_al_20240629.py or 35M-popik_et_al_20240629.py

